# An integrated model for termination of RNA polymerase III transcription

**DOI:** 10.1101/2021.08.27.457903

**Authors:** Juanjuan Xie, Umberto Aiello, Yves Clement, Nouhou Haidara, Mathias Girbig, Jana Schmitzova, Vlad Pena, Christoph W. Müller, Domenico Libri, Odil Porrua

## Abstract

RNA polymerase III (RNAPIII) synthesizes essential and abundant non-coding RNAs such as tRNAs. Controlling RNAPIII span of activity by accurate and efficient termination is a challenging necessity to ensure robust gene expression and to prevent conflicts with other DNA-associated machineries. The mechanism of RNAPIII termination is believed to be simpler than that of other eukaryotic RNA polymerases, solely relying on the recognition of a T-tract in the non-template strand. Here we combine high-resolution genome-wide analyses and *in vitro* transcription termination assays to revisit the mechanism of RNAPIII transcription termination in budding yeast. We show that T-tracts are necessary but not always sufficient for termination and that secondary structures of the nascent RNAs are important auxiliary cis-acting elements. Moreover, we show that the helicase Sen1 plays a key role in a fail-safe termination pathway. Our results provide a comprehensive model illustrating how multiple mechanisms cooperate to ensure efficient RNAPIII transcription termination.

## Introduction

Transcription termination is an essential process that sets the borders between genes, therefore avoiding the interference between neighboring transcription units. Furthermore, transcription termination plays an important role in the maintenance of genome integrity by limiting the possible conflicts between transcribing RNA polymerases (RNAPs) and other cellular machineries involved in DNA replication or repair (reviewed in Porrua and Libri, 2015a).

Transcription termination can be envisioned as a multi-step process consisting in the recruitment of termination factors, the recognition of sequence motifs, RNAP pausing, and finally the release of the RNAP and the transcript from the DNA. This last step involves a remodeling of an intricate network of interactions between the RNAP, the nascent RNA and the DNA template (reviewed in Porrua et al., 2016). Within this network, the interactions between the polymerase and the RNA:DNA hybrid are considered as the main determinant of the stability of the elongation complex (EC) (Kireeva et al., 2000). Most eukaryotic organisms possess three different RNAPs that are specialized in producing different classes of transcripts and seem to adopt different strategies to efficiently terminate transcription. RNAPI is responsible for the synthesis of ribosomal RNAs; RNAPII transcribes all protein-coding genes and several classes of non-coding genes and RNAPIII synthetizes short and abundant transcripts among which all tRNAs, the 5S rRNA, and several additional non-coding RNAs.

The mechanisms of transcription termination of the three polymerases have been extensively characterized in the eukaryotic model *Saccharomyces cerevisiae* and many of the principles uncovered in this organism seem to be highly conserved from yeast to humans (reviewed in Porrua et al., 2016). RNAPI and RNAPII require extrinsic protein factors to terminate transcription. RNAPI pauses when it encounters a Myb-like factor bound to the DNA downstream of each rRNA gene (Merkl et al., 2014; Reiter et al., 2012). The release of the paused RNAPI is then mediated by additional proteins, specifically the Rat1 exonuclease and the helicase Sen1 (El Hage et al., 2008; Kawauchi et al., 2008), which are also major termination factors for RNAPII (see below).

The mechanism of RNAPII transcription termination is more complex and involves the action of a larger number of proteins. There are two major termination pathways for RNAPII (reviewed in Porrua and Libri, 2015a). Transcription termination at protein-coding genes relies on a multi-subunit complex that is responsible for the co-transcriptional cleavage of the pre-mRNA at the poly(A) site and the addition of a poly(A) tail. The downstream portion of the nascent transcript is then targeted by Rat1 (XRN2 in humans), which degrades the RNA molecule until it encounters RNAPII and promotes its release from the DNA (Baejen et al., 2017; Kim et al., 2004; Park et al., 2015; Pearson and Moore, 2013; West et al., 2004).

The second pathway is devoted to termination of non-coding transcription and plays an essential role in the control of pervasive transcription as well as in the biogenesis of snoRNAs (Arndt and Reines, 2015; Porrua and Libri, 2015a). This pathway depends on a complex composed of two RNA-binding proteins, <u>N</u>rd1 and <u>N</u>ab3, and the aforementioned helicase <u>S</u>en1 (i.e. the NNS complex). Whereas Nrd1 and Nab3 recognize specific sequence motifs that are enriched in the target non-coding RNAs, the helicase Sen1 induces the dissociation of the EC (Porrua and Libri, 2013; Porrua et al., 2012; Schulz et al., 2013; Steinmetz et al., 2006; Wlotzka et al., 2011). The mechanisms of action of Sen1 in RNAPII transcription have been extensively characterized at the molecular level by our group and others (Han et al., 2017; Hazelbaker et al., 2013; Leonaitė et al., 2017; Porrua and Libri, 2013; Wang et al., 2019). Briefly, Sen1 uses the energy of ATP hydrolysis to translocate along the nascent RNA towards the transcribing RNAPII and, upon transcriptional pausing, it collides with the polymerase and induces its dissociation from the DNA.

A large body of evidence supports the notion that, in contrast to the other RNAPs, RNAPIII can terminate precisely and efficiently at a particular DNA sequence without the need for accessory proteins (reviewed in Arimbasseri et al., 2013 and Porrua et al., 2016). A typical RNAPIII terminator consists in a stretch of thymidines (T) of variable length in the non-template DNA strand that, according to the current model, is sufficient to promote both pausing and release of RNAPIII. Upon transcription of a T-tract, the weakness of the resulting rU:dA hybrid is thought to be central to the destabilization of the RNAPIII EC (Mishra and Maraia, 2019). The particular sensitivity of RNAPIII to weak rU:dA hybrids compared to other RNAPs that do not sense T-tracts as terminators is believed to depend on the less-extensive interactions between RNAPIII and the RNA:DNA hybrid, (Hoffmann et al., 2015). The Ts in the non-template strand play an additional critical role in transcription termination (Arimbasseri and Maraia, 2015), as they have been proposed to be recognized by the C37 and C53 subunits of RNAPIII that also contribute to termination (Landrieux et al., 2006; Rijal and Maraia, 2013). An alternative model proposed by Nielsen and coauthors (Nielsen et al., 2013) posits that T-tracts are required for RNAPIII pausing but are not sufficient for its release from the DNA.

These authors have proposed that the folding of the nascent RNA into a hairpin-like structure in the vicinity of the paused RNAPIII is an absolute requirement for termination. The hairpin would invade the RNA exit channel of the polymerase, thus provoking its dissociation from the DNA. The proposed mechanism is reminiscent of the so-called intrinsic termination pathway described for bacterial RNAP. This hairpin-dependent model remains, however, highly controversial since it is seemingly in disagreement with a large body of former experimental evidence (Arimbasseri et al., 2014).

The model according to which sequence signals are the sole determinant of RNAPIII termination has also been challenged in the fission yeast *Schizosaccharomyces pombe* by a recent report showing that one of the homologues of the *S. cerevisiae* Sen1 (hereafter designated *Sp* Sen1) is involved in RNAPIII termination *in vivo* (Rivosecchi et al., 2019). Deletion of this gene that in *S. pombe* is non-essential leads to a global shift of RNAPIII occupancy downstream of tRNA genes, consistent with the notion that *Sp* Sen1, in addition to T-tracts, is required for RNAPIII termination in this organism. The precise role of *Sp* Sen1 in termination as well as its mechanism of action were, however, not addressed in this study. Thus, much uncertainty remains about the relative contribution of sequence elements, RNA structures and *trans*-acting factors to the efficiency of RNAPIII transcription termination. Also, to what extent the different termination mechanisms are evolutionary conserved remains an open question.

In the present study we combine high-resolution genome-wide approaches with *in vitro* transcription termination assays using highly-purified components to dissect the mechanism of RNAPIII transcription termination in *S. cerevisiae*. We observe that termination at the primary terminator of RNAPIII-dependent genes (i.e. the first T-tract after the gene), is only partially efficient and, thus, a considerable fraction of polymerases terminate in the downstream region. We provide *in vivo* and *in vitro* evidence that the helicase Sen1 plays a global role in RNAPIII transcription termination and that this function relies on the interaction of its N-terminal domain with RNAPIII. However, we find that Sen1 contributes very little to the efficiency of primary termination and that it mainly functions as a fail-safe mechanism to promote termination of RNAPIIIs that override the first termination signal. Our data indicate that only T-tracts within a particular length range are sufficient to promote autonomous termination by RNAPIII. Nevertheless, we show that tRNA genes often contain suboptimal termination signals and that their capacity to induce termination can be complemented by Sen1 as well as by secondary structures of the nascent RNA. These two factors act in a mutually exclusive manner since the presence of RNA structures prevent the loading of Sen1 onto the transcript, which is strictly required for Sen1-mediated termination. While Sen1 can also promote the release of RNAPIII at pausing sites other than T-tracts, we find that RNA structures can only function in association with canonical termination signals.

Together, our data allow revisiting former models for RNAPIII transcription termination and offer a novel and detailed view of how intrinsic components of the EC (i.e. T-tracts and RNA structures) and the extrinsic factor Sen1 concur to promote efficient termination of RNAPIII transcription.

## Results

### The N-terminal domain of Sen1 interacts with RNAPIII

*S. cerevisiae* Sen1 is a modular protein composed of a large N-terminal domain (aa 1-975), a central helicase domain (aa 1095-1867) and a C-terminal disordered region (aa 1930-2231, see figure 1A). We have recently shown that the N-terminal domain (NTD) is essential for viability and for termination of RNAPII transcription and that it recognizes the CTD of RNAPII, although it is not the only RNAPII-interacting region in Sen1 (Han et al., 2020). In a quest for other functional interactions mediated by the Sen1 Nter, we performed co-immunoprecipitation (co-IP) experiments followed by mass spectrometry (MS) analyses using either a full-length or a *Δ*NTD version of Sen1 as a bait (table 1 and S1). We expressed both *SEN1* variants from the *GAL1* promoter (p*GAL1*) because only overexpression of the *sen1ΔNTD* allele supports viability (Han et al., 2020). In agreement with previous reports (Appanah et al., 2020; Yüce and West, 2013), we detected an RNase-resistant interaction of Sen1 with its partners within the NNS-complex Nrd1 and Nab3, several replication and transcription-related factors, as well as with the three RNAPs. Strikingly, deletion of the NTD abolished the association of Sen1 with RNAPIII and most replication factors without markedly affecting other interactions. Additional co-IP/MS experiments using the isolated NTD as a bait confirmed the interaction with replication factors (e.g. Ctf4) and RNAPIII subunits, strongly suggesting direct protein-protein interactions between the NTD and these factors (table 1 and S2).

**Figure 1:**
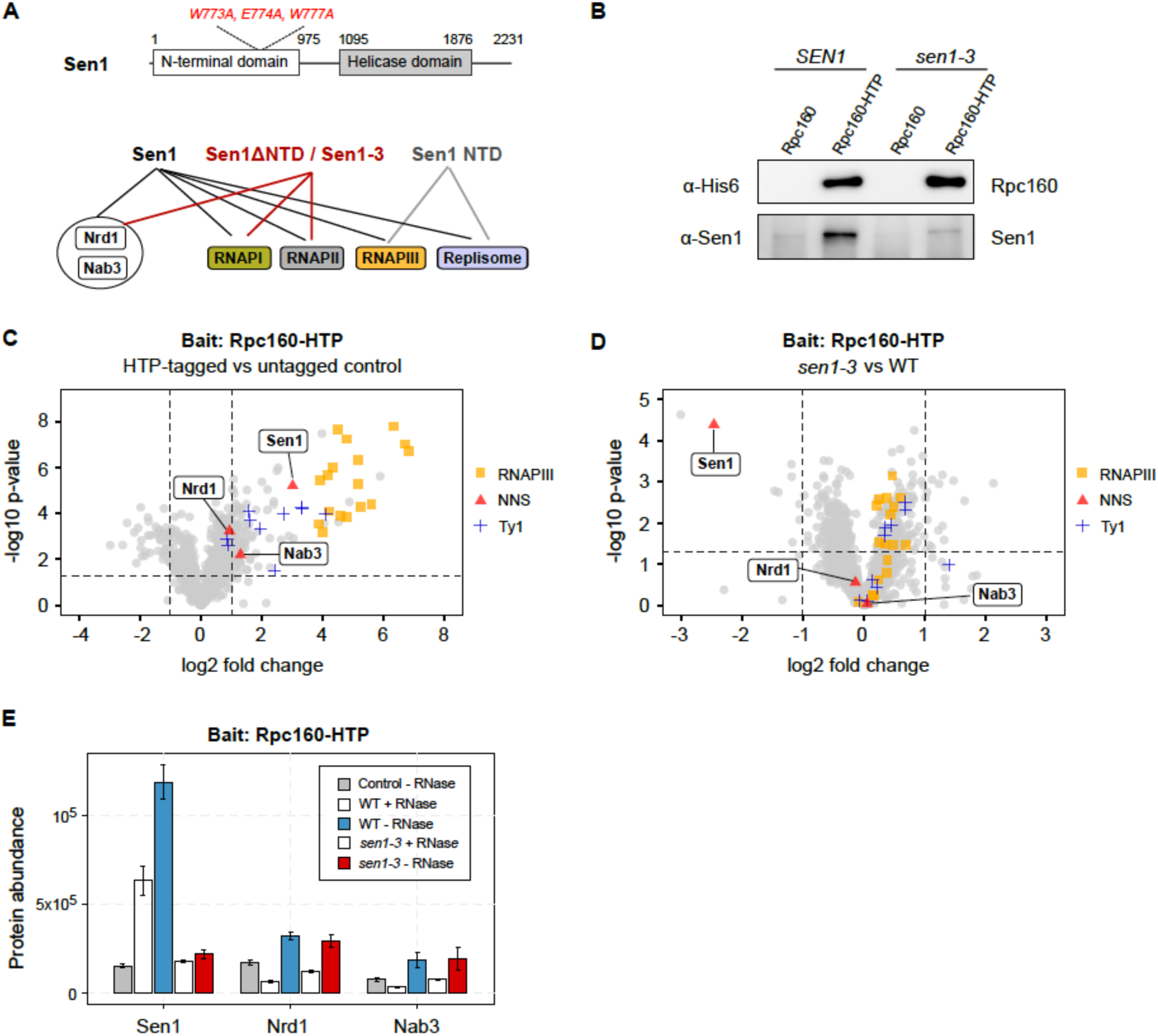
The N-terminal domain of Sen1 interacts with RNAPIII. **A)** Summary of the results of coimmunoprecipitation-MS experiments using different versions of TAP-tagged Sen1 as baits that are included in table 1. A scheme of Sen1 protein indicating the different functional domains as well as the position of the mutations introduced in the *sen1-3* strain is shown on the top. Globular domains are denoted by solid bars while protein regions predicted to be intrinsically disordered are indicated by a line. **B)** Western blot analysis of a representative coimmunoprecipitation experiment using a C-terminally His_6_-TEV-Protein A (HTP)-tagged version of the largest subunit of RNAPIII (Rpc160) as the bait. **C)** and **D)** Label-free quantitative MS analysis of coimmunoprecipitation assays using Rpc160-HTP as the bait. Data correspond to experiments performed in the absence of RNase A treatment. **C)** Volcano plot representing the enrichment of the detected proteins in the HTP-tagged strain relative to the untagged control in a WT (*SEN1*) background. **D)** Quantitative comparison of the proteins that are associated with tagged RNAPIII in a *sen1-3* mutant relative to the WT. Only proteins with a fold change *≥* 2 relative to the control and p-value < 0.05 are considered as significantly changed among the compared conditions. **E)** Comparison of the abundance (arbitrary units) of the different NNS-components in Rpc160-HTP coimmunoprecipitates with or without treatment with RNase A.

**Table 1:** mass spectrometry analyses of coimmunoprecipitation experiments using different versions of Sen1 as bait. Values correspond to the mascot score. ND, not detected. Note that the mascot score depends on the size of the protein and therefore, truncated versions of Sen1 have lower values. ND, not detected. The full datasets and the results of additional replicates are included in tables S1, S2 and S3.

| Protein | Complex | TAP-Sen1 vs TAP-Sen1ΔNter | | | TAP-Nter | | Sen1-TAP vs Sen1-3-TAP | | |
| --- | --- | --- | --- | --- | --- | --- | --- | --- | --- |
| | | Ctrl | Sen1 | Sen1ΔNter | Ctrl | Sen1 Nter | Ctrl | Sen1 | Sen1-3 |
| Sen1 | NNS | 50 | 24819 | 14182 | 153 | 4749 | 0 | 8280 | 7299 |
| Nrd1 | NNS | 0 | 439 | 665 | ND | ND | 0 | 315 | 240 |
| Nab3 | NNS | 0 | 417 | 575 | ND | ND | 0 | 185 | 89 |
| Ctf4 | Replisome | 0 | 2465 | 0 | 19 | 1343 | 0 | 181 | 0 |
| Mrc1 | Replisome | 0 | 40 | 0 | ND | ND | 0 | 63 | 21 |
| Rpa190 | RNAPI | 126 | 463 | 1326 | 31 | 0 | 492 | 2041 | 1812 |
| Rpa135 | RNAPI | 87 | 122 | 664 | 29 | 0 | 400 | 1505 | 1060 |
| Rpb1 | RNAPII | 0 | 2770 | 3003 | 81 | 102 | 113 | 2313 | 2494 |
| Rpb2 | RNAPII | 0 | 2302 | 2328 | 86 | 55 | 34 | 1482 | 1684 |
| Rpc160 | RNAPIII | 0 | 7479 | 0 | 0 | 358 | 25 | 2216 | 59 |
| Rpc128 | RNAPIII | 0 | 4731 | 0 | 0 | 125 | 42 | 938 | 204 |
| Rpc82 | RNAPIII | 0 | 2735 | 0 | 0 | 233 | 0 | 1135 | 34 |
| Rpc53 | RNAPIII | 68 | 1255 | 183 | 0 | 56 | 0 | 500 | 34 |
| Rpc37 | RNAPIII | 0 | 1982 | 0 | 0 | 132 | 0 | 197 | 50 |
| Rpc34 | RNAPIII | 0 | 2022 | 0 | 0 | 28 | 0 | 433 | 174 |
| Rpc31 | RNAPIII | 0 | 1212 | 0 | 0 | 19 | 0 | 426 | 0 |
| Rpc25 | RNAPIII | 0 | 410 | 0 | 0 | 0 | 0 | 79 | 0 |
| Rpc17 | RNAPIII | 0 | 422 | 0 | 0 | 91 | 0 | 128 | 0 |
| Rpc11 | RNAPIII | 0 | 191 | 0 | 15 | 37 | 33 | 138 | 62 |

The interaction of the N-terminal domain of Sen1 with the replisome was found to depend on the replication factors Ctf4 and Mrc1 in a parallel, collaborative study (Appanah et al., 2020). In that work, we found that combination of three point mutations in a conserved region of the Sen1 NTD (W773A, E774A, W777A; defining the Sen1-3 variant) abolishes the interaction with these proteins. Importantly, we showed that Sen1-3 is expressed at similar levels as WT Sen1 and is fully proficient for terminating transcription of NNS target genes (Appanah et al., 2020). To assess whether these mutations also affect the association with RNAPIII, we analysed the protein interactome of Sen1-3 by co-IP/MS (figure 1A, tables 1 and S3). The interaction with RNAPII was not significantly altered in this mutant, in agreement with its proficiency in RNAPII transcription termination (Appanah et al., 2020). Interestingly, we observed that the mutations introduced in Sen1-3 strongly affect the interaction with RNAPIII subunits.

These results are compatible with the notion that the same surface of Sen1 mediates mutually exclusive interactions with the replisome and RNAPIII. Alternatively, the interaction between Sen1 NTD and RNAPIII could be mediated by the replisome. To distinguish between these possibilities, we conducted quantitative MS and western blot analyses on RNAPIII coimmunoprecipitates from WT and *sen1-3* cells (figure 1B-D and table S4). We observed a clear association of RNAPIII with protein components of the Ty1 transposon, which was previously reported and validates our experimental conditions (figure 1C, Bridier-Nahmias et al., 2015). Importantly, while Sen1 was among the most enriched RNAPIII interactors, we did not detect the two replisome anchoring factors, Ctf4 and Mrc1, indicating that Sen1 interacts in a mutually exclusive manner with RNAPIII and the replisome. RNase A treatment induced a ∼2-fold decrease in the level of RNAPIII-bound Sen1, indicating that this interaction is also partially mediated or stabilized by the RNA (figure 1E). As expected, the association of Sen1-3 with RNAPIII was strongly reduced compared to WT Sen1 (figure 1D), even in the absence of RNase treatment, suggesting that the protein-protein interaction mediated by Sen1 NTD is a major pre-requisite for the association of Sen1 with RNAPIII transcripts. Strikingly, the Sen1 NNS partners Nrd1 and Nab3 were very poorly enriched in RNAPIII coimmunoprecipitates (figure 1C-E), strongly suggesting that Sen1 plays a role in RNAPIII transcription termination independently from its function within the NNS-complex.

Taken together, our results support the notion that Sen1 associates with RNAPIII and the replisome within two alternative complexes that are also distinct from the NNS-complex and likely exert different functions.

### Sen1 is required for efficient termination of RNAPIII transcription *in vivo*

The most widely accepted model for RNAPIII transcription termination posits that the polymerases recognize a *cis*-acting element composed of a stretch of thymidines on the non-template DNA and is released without the need for additional *trans*- or *cis-*acting factors (reviewed in Arimbasseri et al., 2013 and Porrua et al., 2016). However, the evidence supporting a direct interaction between RNAPIII and Sen1 prompted us to investigate a possible role for the latter in terminating RNAPIII transcription. To this end, we generated high-resolution maps of transcribing RNAPIII by CRAC (crosslinking analysis of cDNAs) (Candelli et al., 2018; Granneman et al., 2009). Briefly, the nascent RNAs are UV-crosslinked to RNAPIIIs *in vivo* and the RNAPIII-RNA complexes are purified under stringent conditions. The extracted RNAs are then used to generate cDNAs that are deep-sequenced, providing the position of RNAPIIIs with nucleotide resolution. We performed these experiments in WT or *sen1-3* cells as well as in a Sen1-AID (auxin-inducible degron) strain, which allowed assessing the effect of Sen1 depletion (figure 2).

**Figure 2:**
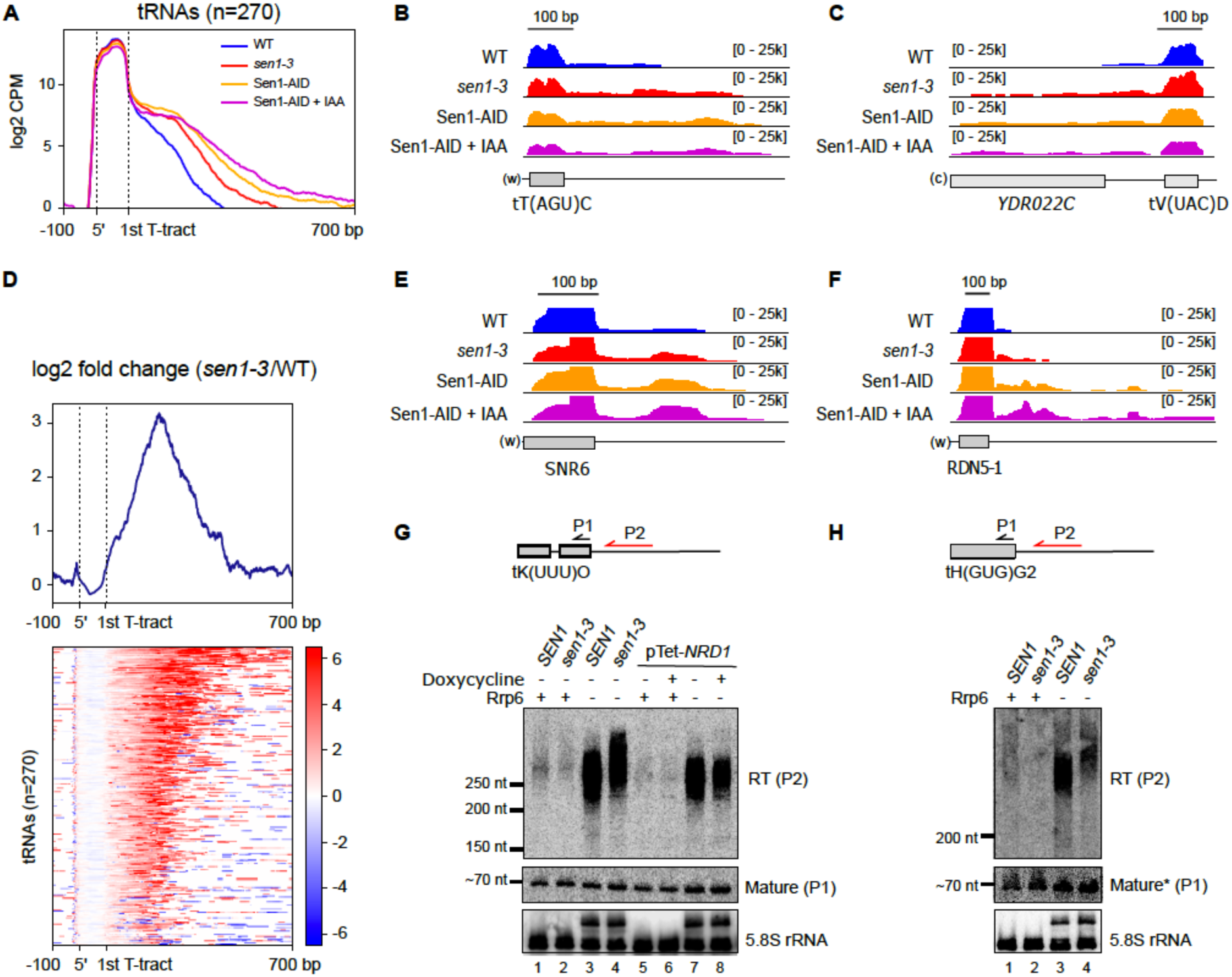
The interaction of Sen1 with RNAPIII is globally required for efficient transcription termination at RNAPIII-dependent genes. **A)** Metagene analysis of the RNAPIII distribution around tRNA genes. The signal covering the region between the 5’ and the primary terminator (i.e. the 1^st^ T-tract after the 3’ end of the mature tRNA) is scaled to 100 bp. Values on the *y*-axis correspond to the mean coverage expressed as counts per million (CPM) multiplied by 10. Sen1-AID denotes the strain expression an Auxin Inducible Degron version of Sen1. IAA: indole-3-acetic acid, an analogue of auxin. **B)** and **C)** Integrative Genomics Viewer (IGV) screenshots of examples of tRNA genes displaying termination defects upon mutation or depletion of Sen1. “w” and “c” denote the Watson and the Crick strands, respectively. The values under brackets correspond to the scale of the RNAPIII signal expressed in 10xCPM. **D)** Heatmap analysis representing the log2 of the fold change (FC) of the RNAPIII signal around tRNA genes in the *sen1-3* mutant relative to the WT. The summary plot on the top was calculated using the average values for each position. **E)** and **F)** Examples of RNAPIII-dependent genes other than tRNA genes that present termination defects upon mutation or depletion of Sen1. **G)** and **H)** Northern blot analysis of transcripts derived from two different tRNA genes in the indicated backgrounds. Schemes on the top indicate the approximate position of the probes (P1 and P2) used for the detection of the different RNA species (RT, for read-through, and mature tRNA). The RNA probe is indicated in red, while DNA oligonucleotide probes are indicated in black (more details in table S5). pTet-*NRD1* denotes strains expressing *NRD1* from a Tet-Off promoter. Depletion of Nrd1 in those strains was achieved by incubation with the tetracycline analogue doxycycline for 10.5h. This system was employed instead of the Nrd1-AID system because RT species are only detectable in a *Δrrp6* background and the Nrd1-AID, *Δrrp6* strain is not viable even in the absence of IAA. The 5.8S rRNA is used as a loading control. Note that Rrp6 is responsible for the processing of the 5.8S rRNA and thus, 5.8S precursors are detected in the *Δrrp6* background. The asterisk in panel H indicates that the signal of mature tH(GUG)G2 corresponds to the same samples loaded in the blot in panel **G**.

We obtained very specific and reproducible RNAPIII occupancy signals in the crosslinked samples relative to un-crosslinked controls, with most reads mapping at RNAPIII-dependent genes (figure S1A-C). Consistent with a former genome-wide study (Turowski et al., 2016), our metagene analyses revealed significant RNAPIII signals downstream of the first T-tract after the 3’ end of tRNA genes, (hereafter referred to as the primary terminator), indicating that termination at this sequence element is only partially efficient *in vivo* (figure 2A-C). Importantly, we observed a clear increase in the RNAPIII signal downstream of the primary terminator in the *sen1-3* mutant, indicating that the interaction with Sen1 promotes termination of RNAPIII, either at the primary terminator or downstream of it. Read-through (RT) transcription was also increased in the Sen1-AID strain even under non-depletion conditions, most likely because the presence of the tag affects the amount or the function of Sen1 even in the absence of auxin as observed for other proteins. Transcriptional read-through was further exacerbated when Sen1 was depleted by the addition of the auxin analogue 3-indoleacetic acid (IAA). The stronger effect of Sen1 depletion relative to the *sen1-3* mutation might imply either that Sen1-3 can still interact weakly with RNAPIII *in vivo*, or that Sen1 functions in RNAPIII termination to some extent in the absence of interaction with the polymerase. Nevertheless, because full depletion of Sen1 also affects termination of many RNAPII non-coding RNA genes, we focused on the more specific *sen1-3* mutant for the rest of our study.

Heatmap analyses of the RNAPIII differential signal (log_2_ ratio) in the *sen1-3* mutant relative to the WT showed that an increase in the signal downstream of the primary terminator could be observed for the vast majority of tRNA genes (figure 2D). Furthermore, inspection of other RNAPIII-dependent genes such as the 5S and U6 genes revealed similar transcription termination defects, indicating that the role of Sen1 in favouring RNAPIII transcription termination is not restricted to tRNA genes (figure 2E-F).

Taken together, our results indicate that Sen1 is globally required for fully efficient termination of RNAPIII transcription *in vivo* and that this Sen1 function relies to a large extent on its interaction with RNAPIII.

### Sen1 functions in RNAPIII transcription independently of the NNS-complex

Nrd1 and Nab3 have been found to bind the precursors of several tRNAs *in vivo* (Wlotzka et al., 2011), and it remain possible that these proteins also partake in RNAPIII termination although they did not appear significantly associated with RNAPIII in our MS analyses (Figure 1E). To address this possibility we conducted RNAPIII CRAC experiments in a Nrd1-AID strain. Depletion of Nrd1 upon treatment with IAA for 1h was sufficiently efficient to provoke clear termination defects at two well-characterized non-coding genes transcribed by RNAPII (i.e. *NEL025c* and *SNR13*, see figure S2C). However, neither the metagene analyses of RNAPIII distribution around tRNAs (figure 3A) nor the inspection of individual RNAPIII-dependent genes (figure 3B-C) revealed any significant effect on RNAPIII transcription termination efficiency. We conclude that, unlike Sen1, Nrd1 is not required for efficient termination of RNAPIII transcription. Because Nab3 is not known to function separately from Nrd1, our results indicate that Sen1 plays a role in RNAPIII transcription independently from the NNS-complex.

**Figure 3:**
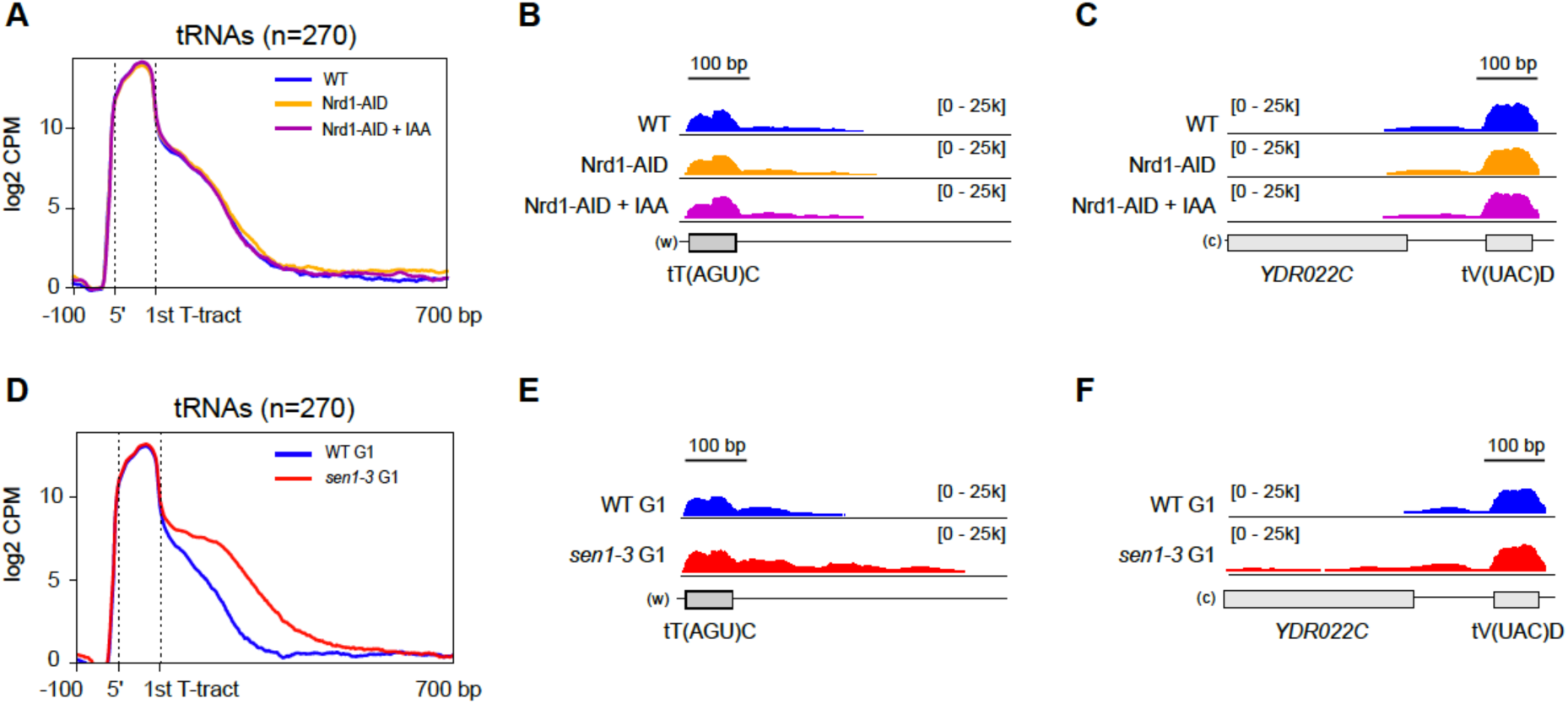
The function of Sen1 in RNAPIII transcription termination does not rely on Sen1 interaction with its partners Nrd1 and Nab3 or with the replisome. **A)** Metagene analysis of the RNAPIII distribution around tRNA genes as in figure 2 in a WT or in a Nrd1 Auxin-Inducible Degron (AID) strain in the absence or in the presence of IAA. Additional experiments validating the efficiency of Nrd1 depletion on well-characterized NNS target RNAs are included in figure S2. **B)** and **C)** Individual examples of tRNA genes that exhibit clear termination defects in a *sen1-3* mutant (see figure 2B-C) but not in Nrd1-depleted conditions. **D)** Metagene analysis as in figure 2A but in cells blocked in the G1 phase of the cell cycle. **E)** and **F)** Individual examples of tRNA genes that display termination defects in the *sen1-3* mutant in cells blocked in G1 as well as in asynchronous cells (compare with figure 2B-C).

### The function of Sen1 in RNAPIII transcription termination is not mediated by the replisome

Our analyses of Sen1 and RNAPIII protein interaction network support a model whereby Sen1 interacts with RNAPIII and the replisome in a mutually exclusive manner. However, they do not exclude the possibility that the replisome mediates the loading of Sen1 onto RNAPIII, for instance when a collision between these complexes occurs (e.g. Sen1 could interact sequentially with the replisome and RNAPIII). RNAPIII transcription units are indeed hot spots of conflicts between the transcription and the replication machineries (Osmundson et al., 2017). Therefore, we considered the possibility that Sen1 might only function in RNAPIII transcription termination in the presence of ongoing replication. To explore this possibility, we performed parallel RNAPIII CRAC experiments in asynchronous cells and in cells arrested in the G1 phase by treatment with *α*-factor, in a WT and a *sen1-3* background (G1-arrest was verified by FACS analysis, figure S1D-E). Importantly, we observed a very similar RNAPIII pattern in G1-arrested and asynchronously growing cells (figure 2A-C and 3D-E), namely prominent RNAPIII termination defects in *sen1-3*.

The finding that abolishing the interaction between Sen1 and RNAPIII reduces the efficiency of termination even in the absence of the replisome (i.e. G1-arrested cells) indicates that Sen1 plays a role in termination of RNAPIII transcription independently of its association with the replisome.

### Sen1 operates in a fail-safe transcription termination pathway

Our genome-wide data indicate that the association of Sen1 with RNAPIII globally increases the efficiency of transcription termination. However, these results are consistent with both a function for Sen1 in promoting termination at the primary terminator and/or a role in removing polymerases that constitutively escape primary termination.

To distinguish between these possibilities, we first analysed the distribution of the RNAPIII CRAC signal in WT and *sen1-3* cells. The total transcription levels, inferred from the RNAPIII signal within the gene body, were virtually identical in WT and *sen1-3* cells, indicating that the mutations in Sen1-3 do not impact transcription initiation or elongation (figure S3A-B).

We then computed for each tRNA gene both the RT index (i.e. the ratio of the RNAPIII signal downstream versus upstream of the primary terminator) and the RT length (i.e. the distance between the primary terminator and the 3’ end of the RT signal) in the WT and in *sen1-3* (figure 4A). For most genes, we observed an increase in the RT index in *sen1-3* cells compared to WT cells (figure 4B-E), which is compatible with Sen1 functioning in primary or in secondary termination, since failure in either one of these processes alone would result in the accumulation of RNAPIIIs within RT regions. However, the heatmap analyses shown in figure 2D revealed that for most tRNA genes, very little or no RNAPIII accumulation could be observed immediately after the primary terminator in the mutant, with the largest increase of RNAPIII signal occurring further downstream, arguing against a major role for Sen1 at the primary termination site. Consistent with this notion, we observed a clear increase in the RT length in the mutant (figure 4B-E), indicating that polymerases that have escaped primary termination transcribe for longer because downstream termination is defective in the Sen1 mutant.

**Figure 4:**
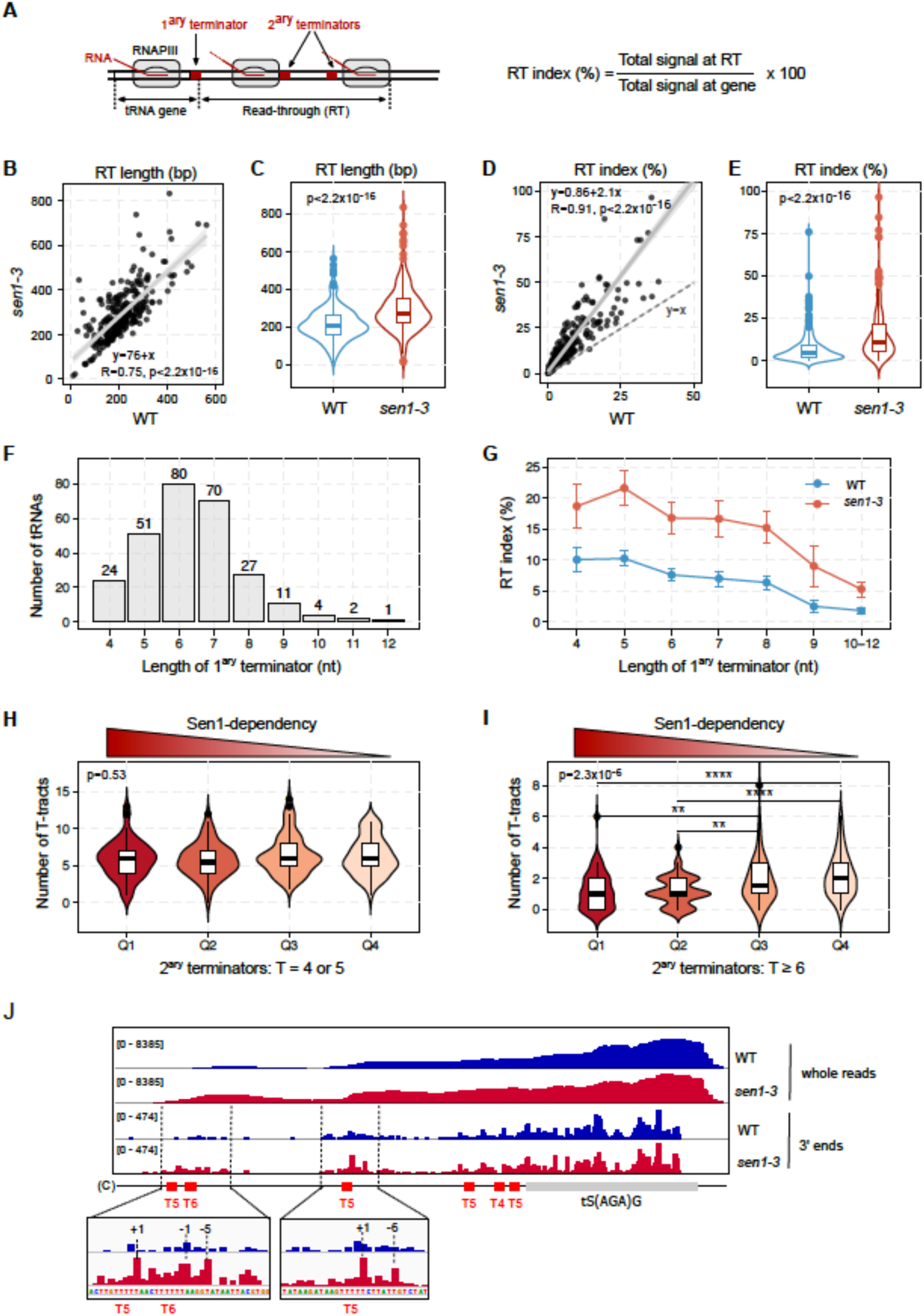
Sen1 functions mainly on secondary termination. **A)** Scheme of tRNA transcription units indicating the relevant elements and parameters used for the assessment of the transcription termination efficiency in the WT and the *sen1-3* mutant. **B)** and **C)** Comparison the RT length for the different tRNA genes in the mutant relative to the WT. **B)** Correlation plot. The grey zone corresponds to the confidence interval whereas R is the Pearson’s correlation coefficient. *p* is the p-value associated with Pearson’s correlation. **C)** Violin plot showing the distribution of RT lengths in the WT and in sen1-3. The p-value (*p*) was calculated with the Wilcoxon test. **D)** and **E)** Comparison of the RT index measured in the indicated strains for each tRNA gene. **D)** Correlation plot generated as in **B)**. Three outliers in *sen1-3* are not shown. E) Violin plots as in **C)** but with RT index values. Three outliers in *sen1-3* are excluded. Note that both RT length and the RT index are inversely proportional to the termination efficiency (e.g. higher RT index indicates lower termination efficiency). **F)** Histogram representing the number of tRNA genes that possess a primary terminator of each indicated length. Only consecutive thymidines are considered when computing the length of the primary terminator. **G)** Analysis of the RT index of tRNA genes grouped according to the length of their primary terminator in either the WT or the *sen1-3* mutant. Data points correspond to the average value whereas error bars denote the standard error. **H)** and **I)** Analysis of the number of either “weak” (H) or “strong” (I) terminators located at the 700 bp region downstream of the primary terminator for tRNA genes grouped according to the extent of termination defects in the *sen1-3* mutant (i.e. dependency on Sen1 for efficient transcription termination). Groups correspond to quartiles (Q) defined by the tRNA gene ranking obtained in the heatmap analyses in figure 2D, where Q1 includes the 25% of genes with the highest impairment in transcription termination in the *sen1-3* mutant. *p* corresponds to the p-value for the global comparison of the four groups according to the Kruskal-Wallis test. Asterisks denote the p-values of pairwise comparisons (*: p ≤ 0.05 ; **: p ≤ 0.01; ***: p ≤ 0.001; ****: p ≤ 0.0001). **J)** IGV screenshot of an individual tRNA showing the distribution of RNAPIII CRAC signal in the WT and the *sen1-3* mutant. The 3’ ends datasets provide the position of individual RNAPIIIs with single-nucleotide resolution. Insets are zoom in views of the main regions where RNAPIII accumulates in the mutant. Coordinates in insets correspond to the position relative to the beginning of the nearest downstream T-tract.

Because termination defects would lead to the production of different RNA species from tRNA genes depending on whether they occur at the primary terminator or at read-through regions, we set out to analyse these RNAs by northern blot (figure 2G-H). Mature tRNAs are generated by termination at the primary terminator and eventually by the processing of short 5’ and 3’- extentions. Therefore, defects in primary termination are expected to result in lower amounts of mature tRNAs with a concomitant increase in the amount of RT transcripts. We could only detect RT RNAs for the tRNA genes tK(UUU)O and tH(GUG)G2 in the absence of the exosome-associated exonuclease *RRP6* (figure 2G-H), consistent with former data indicating that RT species are degraded by the RNA exosome (Turowski et al., 2016). In the case of tG(GCC)F2, simultaneous deletion of *RRP6* and depletion of the tRNase Z endonuclease Trz1, involved in the processing of tRNA precursors (Skowronek et al., 2014), was required for the strong detection of RT transcripts (figure S2A), indicating that RT transcripts can also be targeted by Trz1.

Importantly, in all these cases we did not observe a significant decrease in the abundance of mature tRNAs in *sen1-*3, not even upon depletion of Trz1, excluding the possibility that RT transcripts are recognized as tRNA precursors by this endonuclease and cleaved to generate mature tRNAs (figure S2A and data not shown). Accordingly, the overall abundance of RT RNAs was similar in the WT and in *sen1-*3, but these species were globally longer in *sen1-3* cells, confirming CRAC data suggesting that they result from defective Sen1-dependent termination occurring downstream of tRNA primary terminators (figures 2G-H and S2A). This increase in size was not observed, as expected, when the NNS subunit Nrd1 was depleted, consistent with the Nrd1-AID RNAPIII CRAC data (figures 2G and 3A-C).

To further support the notion that Sen1 functions mainly on RNAPIIIs that have escaped the primary termination site, we performed more detailed analyses of our CRAC data.

If Sen1 does not function in primary termination, its failure to interact with RNAPIII should affect similarly genes with weak or strong primary terminators. Based on *in vitro* data, the minimal length for a functional terminator is 5 Ts (Arimbasseri and Maraia, 2015; Mishra and Maraia, 2019) but 6 Ts are required for relatively efficient termination and it is generally assumed that the termination efficiency is higher as the T-tract length increases. In partial agreement with these notions, we observed that i) the first T-tract rarely contains 4 Ts, ii) 6 Ts and 7 Ts are the most frequent terminators at this position and iii) tracts longer than 8 Ts are rarely found as primary terminators (figure 4F). We analysed the RT index of tRNAs clustered according to the length of their primary terminator and, as expected, we found that the RT index in these clusters tends to decrease as the T-tract length increases (figure 4G) in inverse correlation with the termination efficiency. Importantly, in *sen1-3* cells the RT index increases similarly for all clusters suggesting that having an inefficient primary terminator does not make termination more sensitive to Sen1, arguing against a role of Sen1 at these sites.

The region downstream of tRNA genes contains T-stretches that were previously proposed to play a role as secondary termination sites (Turowski et al., 2016). We considered the possibility that Sen1 might be preferentially required for tRNA genes having a lower number of secondary termination sites or less efficient ones. To address this possibility, we ranked the different tRNAs according to the extent of the RNAPIII accumulation in *sen1-3* relative to WT cells, thus defining a hierarchy of Sen1-dependency. For each tRNA, we computed the number of weak (4 or 5 Ts) or strong (≥ 6 Ts) terminators in regions of secondary termination and we compared the average number of terminators of each kind in the different quartiles. Interestingly, we found that the tRNA genes that are more dependent on Sen1 for termination (i.e. Q1) tend to have a lower number of efficient terminators compared to those that are less dependent (i.e. Q3 and Q4). In contrast, the number of weak terminators, which have a lower impact on RNAPIII progression, was similar in all groups of tRNAs (figures 4H-I and S3).

These results strongly suggest that Sen1 compensates for the lack of efficient terminators in regions of secondary termination. This could imply that Sen1 improves termination at weak terminators or that it promotes termination at other sequences. A careful analysis of the RNAPIII CRAC signal at individual tRNA genes provided evidence supporting both possibilities (figure 4J and S4). Mapping only the 3’ end of the nascent RNA allows obtaining a precise readout of RNAPIII position with single-nucleotide resolution. We observed very little if any effect of the *sen1-3* mutation at positions around the primary terminator, while RNAPIII was clearly found to accumulate preferentially around T-tracts, but also at other sequences, in the downstream regions. Together our results support the notion that Sen1 does not play a prominent role in primary termination and rather promotes the release of RNAPIIIs that pause within regions of secondary termination.

### Sen1 can promote termination of RNAPIII transcription *in vitro*

We have previously demonstrated that Sen1 can directly promote termination of RNAPII transcription in a sequence-independent manner (Porrua and Libri, 2013). To assess whether Sen1 can also directly induce RNAPIII transcription termination and whether it requires the presence of canonical termination signals, we employed an *in vitro* transcription termination system containing purified proteins (i.e. RNAPIII and full-length Sen1), transcription templates and nascent RNA (figures 5A and B).

**Figure 5:**
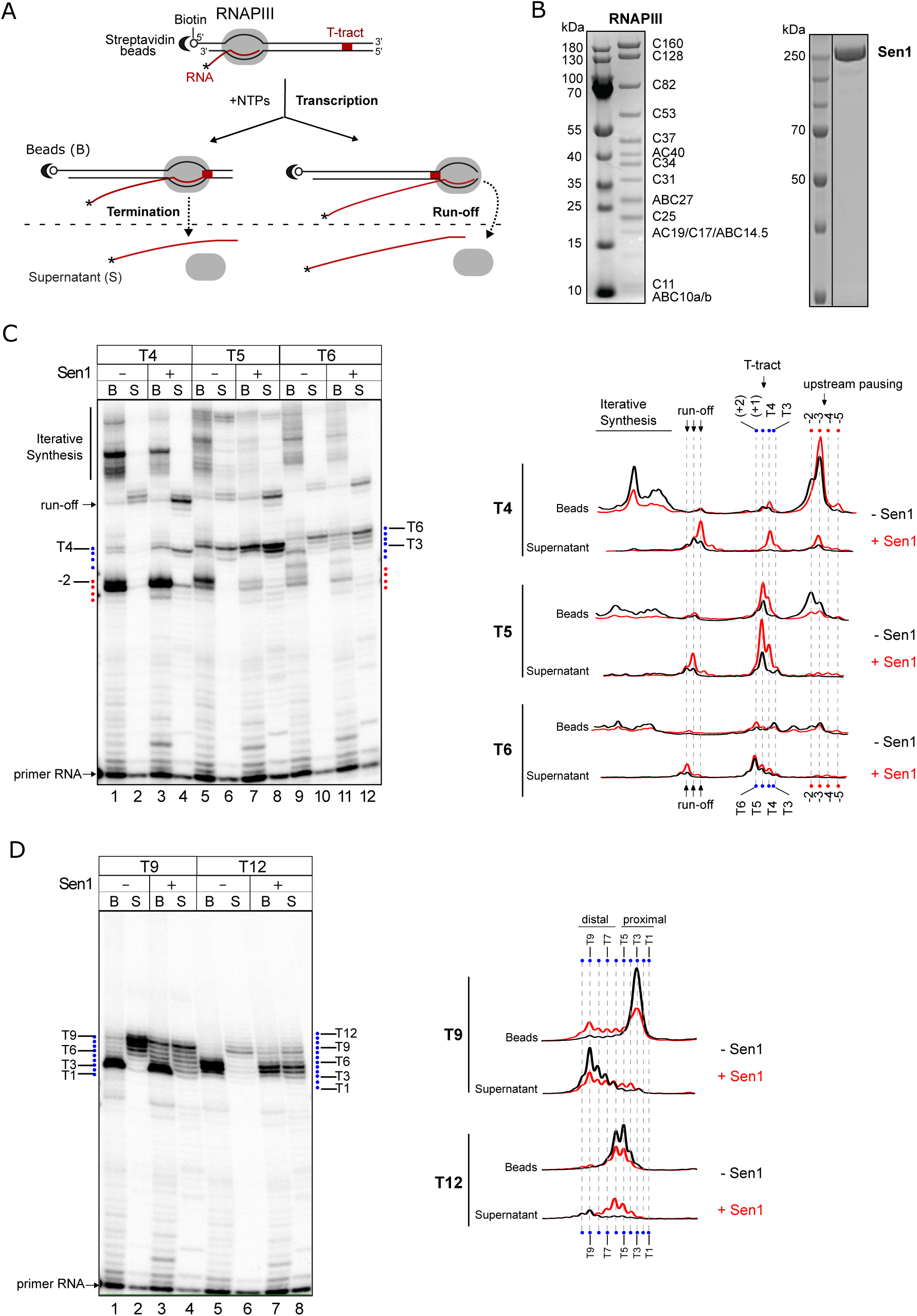
Sen1 can induce termination of RNAPIII transcription *in vitro*. **A)** Scheme of an *in vitro* transcription termination (IVTT) assay. Ternary elongation complexes (ECs) composed of purified RNAPIII, nascent RNA, and DNA transcription templates are assembled by step-wise addition of the different components (see methods) and associated with streptavidin beads via the 5ʹ biotin of the non-template strand to allow subsequent separation of beads-associated (B) and supernatant (S) fractions. The RNA is radioactively labeled at the 5’end to allow its detection (indicated by an asterisk). Each transcription template contains a T-tract of a particular length on the non-template strand. After addition of nucleotides, RNAPIII transcribes and can pause at different positions, including the T-tract. RNAPIIIs that pause at a T-tract can either dissociate from the DNA to the supernatant (i.e. undergo transcription termination) or remain paused, and thus associated with the beads, or resume transcription. Polymerases that read-through the T-tract and reach the end of the template can either run-off, with concomitant release of full-length transcripts into the supernatant, or perform iterative synthesis. For transcripts associated with paused RNAPIIIs, the comparison of the fraction that is retained in the beads with the fraction that is released provides an estimate of the efficiency of termination at each site. **B)** SDS-PAGE analyses of the protein preparations used in IVTT assays. **C)** Analyses performed on templates containing T-tracts composed of 4 (T4), 5 (T5) or 6 (T6) consecutive Ts. Left: Denaturing PAGE analysis of transcripts from an IVTT assay testing the capacity of Sen1 and T-tracts of different lengths to induce RNAPIII transcription termination. “B” corresponds to the beads fraction while “S” denotes the supernatant. Representative gel of one out of three independent experiments. Right: Profile of the signal over the region of interest for each gel lane. **D)** Analysis of IVTT reactions performed on templates containing stretches of 9 (T9) or 12 (T12) Ts. These reactions were performed in parallel with those in panel **C)** but migrated on different gels. Left: Representative gel of one out of three independent experiments. Right: Profile of the signal over the region of interest for each gel lane. The position of the nucleotides of interest was determined by migrating in parallel a radioactively-labelled ladder (not shown).

We first analysed the capacity of canonical terminator sequences to induce RNAPIII transcription termination by comparing the behaviour of RNAPIII on transcription templates containing T-tracts of variable lengths (i.e. from 4 to 12 Ts, see figures 5C-D and S5). Consistent with former data (Arimbasseri and Maraia, 2015; Mishra and Maraia, 2019), we observed only very weak polymerase pausing at the T4 terminator and no detectable RNAPIII release. The T5 terminator induced stronger pausing and intermediate levels of RNAPIII release, while the T6 terminator promoted very efficient release. Stretches of 9 or 12 Ts induced very strong RNAPIII pausing as virtually no transcription signal could be detected downstream of these terminators but a substantial proportion of RNAPIIIs remained associated with the proximal part of these long T-tracts (∼ 50% for the T9 and ∼ 80% for the T12). This might be due to the recognition of the distal portion of the T-tract in the downstream DNA by RNAPIII, which might induce strong pausing and disfavour release (see Discussion). Indeed, for shorter T-tracts we also observed several prominent pausing sites a few nt upstream of these sequences, both *in vitro* (figure 5C) and *in vivo* (figures 4J and S5) supporting the idea that RNAPIII can sense downstream untranscribed T-tracts.

Importantly, the presence of Sen1 in the reaction provoked a substantial increase in the levels of transcription termination at the T4 terminator, and, to a lesser extent, at the T5 terminator, while no significant effect was observed for the more efficient T6 terminator. This result indicates that Sen1 can enhance RNAPIII release at weak terminators.

Interestingly, we found that Sen1 could also promote the release of RNAPIIIs that are paused at the proximal part of long T-tracts, especially in the case of the T12 terminator, for which roughly 50% of paused RNAPIIIs were released by Sen1. Finally, and importantly, we also observed Sen1-dependent release of RNAPIIIs that are paused at sequences other than T-tracts, for instance, at pausing sites upstream of the canonical terminators, which corroborates our *in vivo* RNAPIII CRAC analyses (figure 4J and S4). Taken together, these results indicate that Sen1 can both enhance termination at inefficient terminators and promote termination at unrelated sequences.

### Sen1 employs a similar mechanism to terminate transcription of RNAPII and RNAPIII

According to previous studies, canonical terminators contain signals that induce both RNAPIII pausing and release from the DNA (reviewed in Arimbasseri et al., 2013 and Porrua et al., 2016). The above results indicate that Sen1 requires polymerase pausing but not necessarily the presence of a T-tract for terminating RNAPIII. To further explore this idea, we performed *in vitro* transcription assays with modified templates containing a G-less cassette followed by a run of Gs to force the stalling of RNAPIII at the G-stretch in the absence of GTP (figure 6A). In these conditions, and similarly to what was observed for RNAPII, Sen1 could induce the release of roughly 50% of paused RNAPIIIs, demonstrating that it can terminate transcription at pausing sites other than T-tracts.

**Figure 6:**
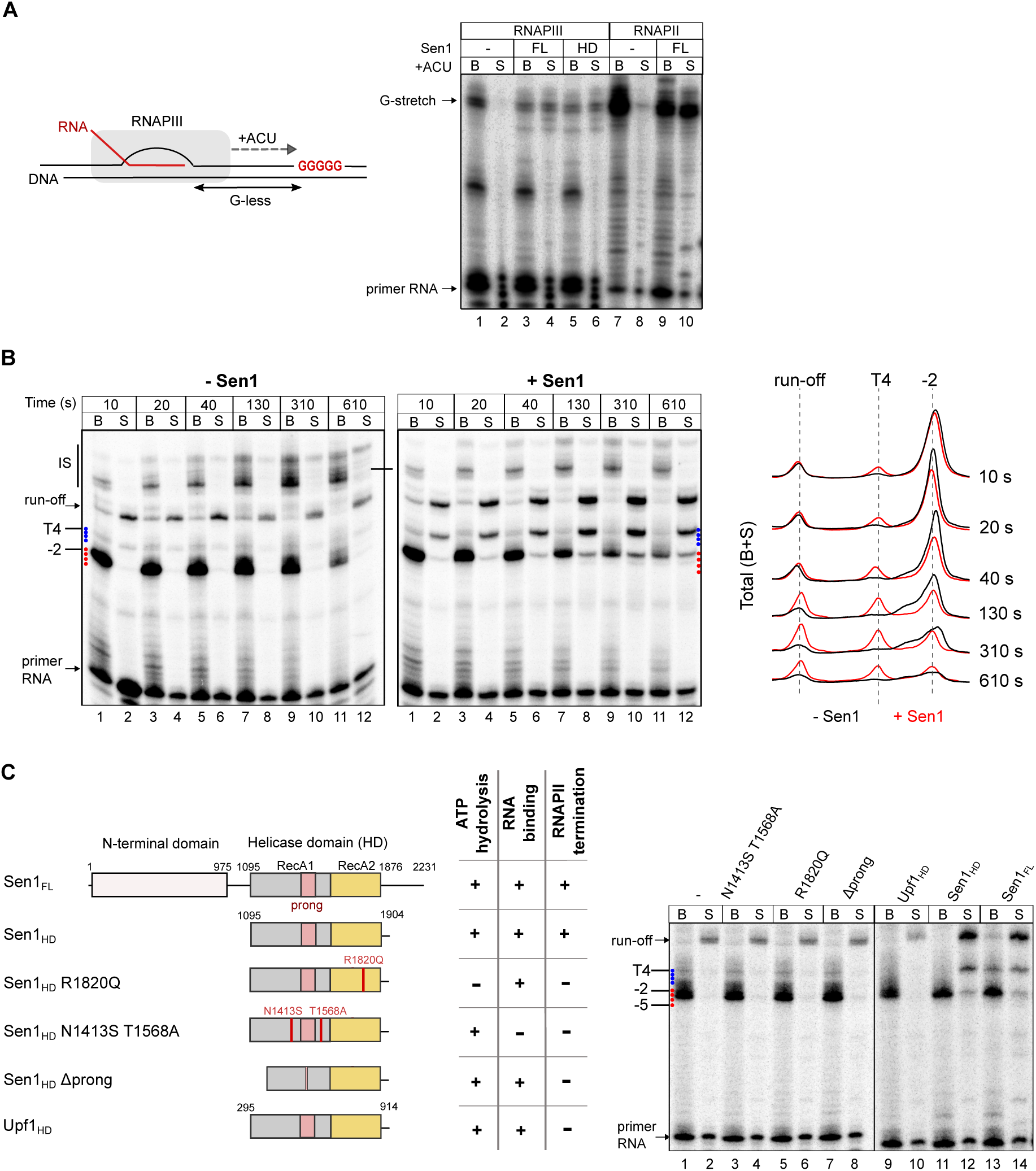
Analysis of the mechanisms of Sen1-mediated termination of RNAPIII transcription. **A)** Analysis of the capacity of Sen1 to promote the release of RNAPIII paused at a sequence other than a T-tract. The transcription templates contain a G-less cassette followed by a stretch of Gs so that in the absence of G in the nucleotide mixture RNAPs are stalled at the first G. Experiments were performed in parallel with purified RNAPII as a positive control for Sen1 activity. The efficiency of transcription termination observed for RNAPII is similar as in our former studies (Han et al., 2017; Leonaitė et al., 2017). **B)** Time-course IVTT assay performed on transcription templates containing a T4 terminator. All the reactions were performed in parallel but were migrated on different gels. Left: Representative gel of one out of two independent experiments. Right: Profile of the total signal (beads and supernatant) over the region of interest. **C)** Analysis of the role of different protein regions and activities of Sen1 in RNAPIII transcription termination. Left: Scheme of the different proteins used in IVTT assays and summary of the relevant phenotypes. The different variants of Sen1 helicase domain (HD) were purified and characterized within the frame of a previous study (Leonaitė et al., 2017). A constitutively active version of the Upf1 HD that possess helicase activity but cannot induce RNAPII transcription termination *in vitro* is used as a negative control (see Porrua and Libri, 2013). The *Δ*prong version of Sen1 HD contains a deletion of amino acids 1461-1554, which corresponds to most of this subdomain. Right: Representative gel of one out of two independent experiments. All the reactions were performed in parallel but were migrated on different gels.

We next set out to investigate the mechanism by which Sen1 induces RNAPIII transcription termination. We have previously shown that in order to dissociate the RNAPII elongation complex, Sen1 needs to load on the nascent RNA and translocate towards the RNAPII using the energy of ATP hydrolysis. Upon colliding with RNAPII, Sen1 can induce forward translocation of the polymerase, which, in the appropriate context, results in its release from the DNA (Han et al., 2017). Our *in vitro* assays support a similar mechanism for RNAPIII release (figure 5C), with evidence for “pushing” the elongation complex at sites of pausing. This is for instance manifest at the T5 terminator (figure 5C, compare lanes 5-6 to 7-8) where a decrease in the pausing signal at position -2 is not due to release at this position but rather by forward translocation and release at a downstream site. This is best illustrated in a time-course experiment performed with the T4 template, in which we quantified the total signal in the presence and absence of Sen1 (figure 6B, right). If a decrease in the signal from a paused polymerase is due to its release, the total signal (i.e. beads + supernatant) at that position should not change. On the contrary, if polymerases are “pushed” by Sen1 and eventually released at a later stage the signal distribution should be shifted downward. Indeed, upon addition of Sen1 we observe such a signal shift as well as the accumulation of RNA signal over time at positions where Sen1 induces its release. These findings support the notion that Sen1 promotes both RNAPIII translocation and its release from the DNA, similarly to what we previously showed for RNAPII.

To further explore the mechanisms of RNAPIII termination by Sen1, we first assessed whether the interaction of Sen1 with RNAPIII, mediated by its N-terminal domain, is required for the actual step of polymerase release. To this end we first analysed the capacity of the helicase domain of Sen1 alone to induce termination *in vitro*. We have previously shown that this domain is sufficient for inducing termination of RNAPII transcription (Han et al, 2017, Leonaite et al, 2017). Strikingly, we found that the helicase domain of Sen1 could induce termination of RNAPIII transcription *in vitro* as efficiently as the full-length protein (figure 6C), suggesting that the association of Sen1 with RNAPIII via its N-terminal domain is not a strict requirement for termination but might rather play a role in the recruitment of Sen1 to RNAPIII *in vivo*. As a negative control, we assessed a catalytically active version of the closely-related helicase Upf1 (Chakrabarti et al., 2011), which could not provoke termination of RNAPIII transcription, indicating that termination is not induced unspecifically by any active helicase but rather requires specific Sen1 activities or features (figure 6C). Finally, we analysed several mutant variants of the Sen1 helicase domain that are deficient for RNA binding (N1413S T1568A) or ATP hydrolysis (R1820Q), or a mutant that retains the catalytic activity but lacks the “prong” domain (i.e. *Δ*1461-1554), which is essential for viability and for RNAPII transcription termination (Leonaitė et al., 2017). Importantly, none of these mutants could promote RNAPIII transcription termination *in vitro*, indicating that Sen1 employs the same structural features and activities to induce transcription termination of RNAPII and RNAPIII.

### RNA structures upstream of T-tracts can promote the release of paused RNAPIIIs

The above results indicate that, akin to the RNAPII system, Sen1-mediated termination of RNAPIII transcription involves Sen1 translocation along the nascent transcript, and our former structural and biochemical data showed that Sen1 can only interact with single-stranded RNA (Porrua and Libri, 2013, Leonaite et al, 2017). tRNAs are highly structured RNA molecules and for a vast majority of them (i.e. 251 out of 270 tRNAs) the spacer between the 3’ end of the mature tRNA and the primary terminator is at most 7 nt. We envisioned that a possible reason for which Sen1 does not function at sites of primary termination is that its binding to the nascent RNA is hindered by the co-transcriptional formation of stable structures in the vicinity of the primary terminator. Conversely, less structured RNAs in the read-through region would allow Sen1 loading and function.

To explore these possibilities we performed *in vitro* transcription assays with modified transcription templates containing a natural hairpin from the 5S RNA, an RNAPIII-dependent transcript, upstream of T-tracts of different lengths. As a control we used a mutated hairpin with substitutions in the left arm preventing stem formation (figure 7A-C and S5). Surprisingly, the presence of a hairpin in the transcribed RNA could significantly stimulate transcription termination at a T4 terminator, similarly to the addition of Sen1 to the unstructured version of the same RNA (figure 7A).

**Figure 7:**
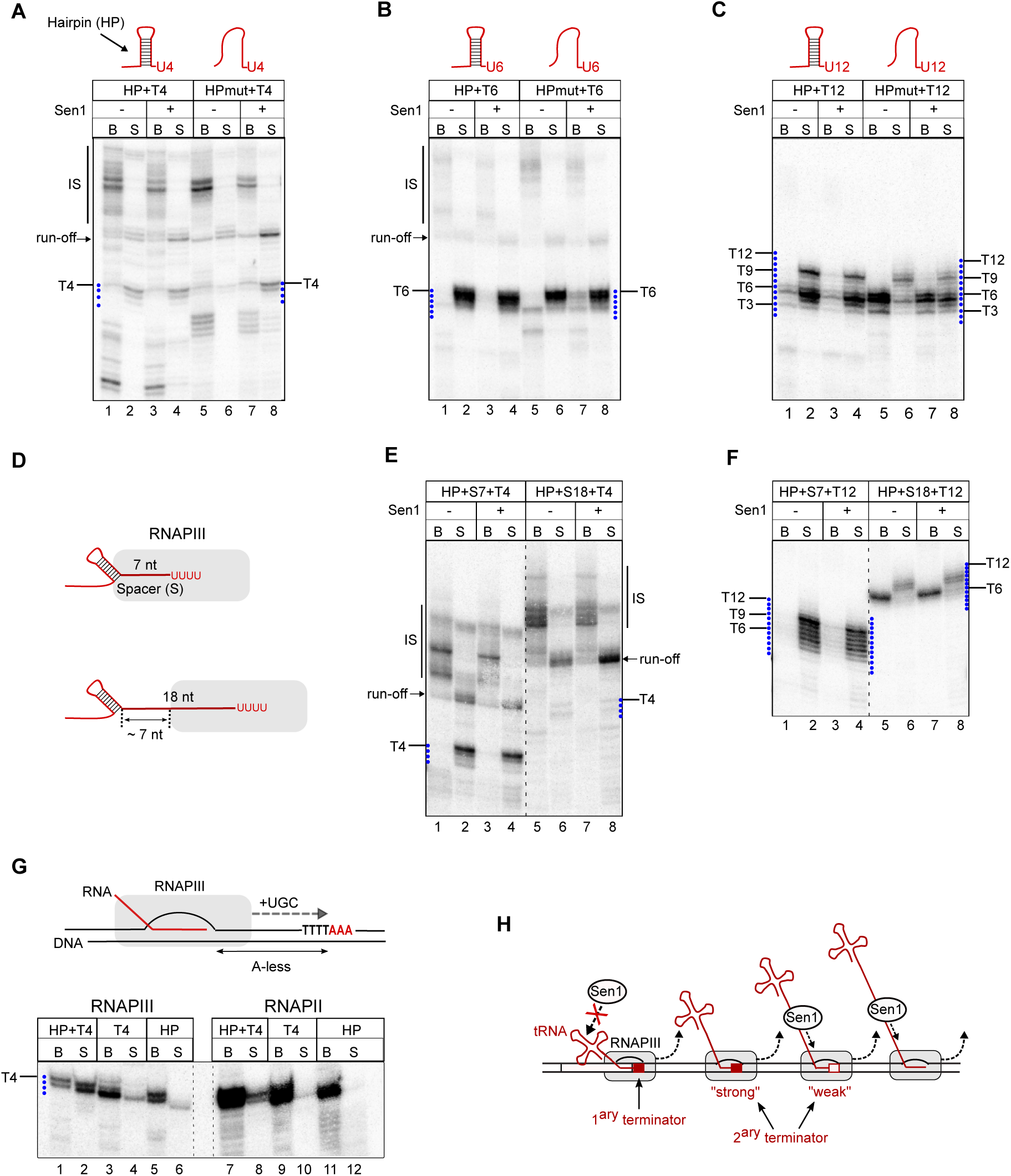
Hairpin-like structures forming in the nascent RNA can complement the function of canonical termination signals. **A-C)** Analysis of the role of RNA structures in transcription termination at a T4 (**A**), T6 (**B**) or a T12 (**C**) terminator. IVTT assays as in figure 5 but with modified templates to introduce a hairpin (HP) in the transcribed RNA immediately upstream of the T-tract. The control template (HPmut) harbors several mutations at the left arm that disrupt hairpin formation (see figure S4 for sequence details and structure prediction of the resulting RNAs). **D-F)** Analysis of the impact of the positioning of the hairpin relative to the T-tract on its capacity to stimulate RNAPIII release. Experiments performed as in **A)** and **C)** but with modified templates to introduce a spacer (S) of the indicated lengths between the hairpin and the T-tract (see figure S4 for details). **D)** Scheme showing the predicted position of the hairpin relative to RNAPIII in the presence of the indicated spacers. IVTT assays with templates containing either a T4 terminator **(E)** or a T12 terminator **(F)**. **G)** Functional dissection of a transcription terminator composed of an RNA hairpin and a stretch of 4 Ts (T4). IVTT assays performed as in **A)** but with modified templates (see figure S4 for details) to include an A-less cassette followed by a stretch of As so that, in the absence of A in the nucleotide mixture, RNAPs are stalled at the first A. In the HP template the T-tract is mutated to CTCT. Experiments were performed in parallel with purified RNAPII to compare the sensitivity of both RNAPs to termination signals. **H)** Model for the role of canonical termination signals, RNA structures and Sen1 in termination of RNAPIII transcription (shown for tRNA genes). At the primary terminator, termination typically involves the action of a T-tract and the secondary structure of the nascent tRNA. RNA structures are required only for T-tracts of sub-optimal length. In downstream regions transcription by RNAPIII is typically terminated either by “strong” secondary terminators, without the aid of Sen1, or by “weak” termination signals if Sen1 can access and load onto the nascent RNA. Sen1 can in principle also promote termination at sites of pausing other than T-tracts.

This result indicates that not only Sen1 but also RNA secondary structures can improve the function of weak terminators. In agreement with this idea, the presence of the hairpin did not enhance termination at the T6 terminator, since this sequence already supports full release of paused RNAPIIIs (figure 7B). However, the RNA structure could induce the release of polymerases paused at the proximal part of the T12 terminator, even more efficiently than Sen1 (figure 7C). These observations support the notion that, similarly to Sen1, RNA hairpins have the capacity to promote RNAPIII release.

Because RNA structures naturally form close to the primary terminator of RNAPIII-dependent genes, we next assessed to what extent secondary structures need to be in proximity to T-stretches to function in termination. To this end, we compared the efficiency of termination on templates containing a T4 or a T12 terminator when the hairpin was located immediately upstream (figure 7A-C), 7 nt or 18 nt upstream of the corresponding T-tract (figure 7D-F). We observed a clear enhancement of RNAPIII release at both T4- and T12-containing templates when the hairpin was located immediately upstream or 7 nt away from the T-tract but not in the presence of a 18 nt spacer. These results indicate that RNA structures can enhance transcription termination only when they are in close proximity to T-tracts. The results with the 18 nt spacer provided a tool to address the impact of secondary structures of the RNA on the function of Sen1 in termination. Indeed, in this case, formation of a hairpin 18 nt upstream of the T-tract only allows exposing a segment of roughly 7 nt of single strand RNA outside of the RNAPIII, which is not sufficient for the loading of Sen1 (Leonaite et al, 2017). Importantly, Sen1 could not release RNAPIII at the T4 terminator in this construct, likely because of the insufficient single stranded RNA span between the structure and the polymerase. We observed, however, an increase in the amount of released run-off transcripts in the presence of Sen1, indicating that at further downstream positions Sen1 can load on the nascent RNA and promote the release of polymerases at the end of the template (figure 7E, lanes 5-8).

Taken together, our results strongly suggest that RNA secondary structures forming in the vicinity of weak primary terminators can markedly improve their function. However, they can also hamper the recruitment of Sen1 to the nascent RNA and, thus, would likely prevent Sen1 from functioning at primary terminators, regardless of their strength.

### RNA secondary structures can form within RNAPIII

A previously proposed model (Nielsen et al., 2013) posits that T-tracts are sufficient for RNAPIII pausing but not for its release from the DNA, for which RNA secondary structures would be strictly required. This model opposes the most widely accepted one, which points to an exclusive role for the T-tract in termination. Our data indicate that secondary structures can promote RNAPIII release but only at defective terminators. We decided to further investigate the mechanism of action of RNA structures in termination. The model of Nielsen et al. postulates that one of the main functions of T-tracts would be to promote RNAPIII backtracking to bring the polymerase in contact with the nearest upstream structure, which would invade the RNA exit channel of RNAPIII, thus destabilizing the elongation complex. Nonetheless, we have observed that the RNA hairpin is functional when located immediately upstream or very close to a T4 or a T12 sequence, which implies either that i) RNAPIII transcribes beyond the T-tract to allow formation of the hairpin, pauses at downstream sequences and undergoes subsequent backtracking; or that ii) the RNA folds at least partially within the polymerase to induce its release. The former possibility appears to be hardly compatible with the case of the T12 terminator, for which we did not find any evidence of RNAPIII transcribing through the terminator (figure 5D).

To address these possibilities, we conducted *in vitro* transcription assays with modified templates where the RNA hairpin is encoded in an A-less cassette and is followed by a T4 sequence and three As (figure 7G and S5). By performing reactions with these templates in the absence of ATP, polymerases cannot transcribe through the terminator and stall at the fourth T of the T4. In these conditions, the hairpin cannot form outside of the polymerase, its downstream arm still being within the polymerase.

Interestingly, we observed that stalled RNAPIIIs were released in the presence of the hairpin but not the corresponding mutant version, indicating that transcription through the terminator is not required for the folding of the hairpin, which must occur in the RNA exit channel of RNAPIII. Importantly, very little, if any, polymerase release was observed when the T4 sequence was mutated, even in the presence of the hairpin, indicating that the hairpin is not an autonomous termination signal and can only function together with a canonical termination sequence. Finally, the concomitant presence of an RNA hairpin and a T-tract induced poor release of RNAPIIs, indicating that these nucleic acid elements function specifically as RNAPIII termination signals.

These findings strongly support the notion that the RNA can fold within the RNAPIII and that backtracking is not required to promote RNAPIII termination. Together, our data comfort the notion that RNA secondary structures are not absolutely required for RNAPIII termination, but can nevertheless function as auxiliary elements that work in concert with weak or defective termination signals.

## DISCUSSION

RNAPIII synthetises short ncRNAs like tRNAs and the 5S rRNA that are absolutely essential for mRNA translation and, therefore, for cell growth and survival. Timely termination of RNAPIII transcription is critical not only for the correct synthesis of these RNAs but also for preventing RNAPIIIs from invading neighbouring genomic regions and interfering with the function of other machineries that operate in these regions. This is even more relevant considering the high expression levels of RNAPIII-dependent genes.

The traditional model for termination of RNAPIII transcription posits that termination exclusively relies on the presence of T-tracts at the 3’ end of RNAPIII-dependent genes that are specifically recognized by the polymerase as termination signals. The implication of additional *cis*-acting factors, such as RNA secondary structures has been previously proposed (Nielsen et al., 2013) but has remained hitherto controversial (Arimbasseri et al., 2014).

Here we show that the mechanisms governing RNAPIII transcription termination *in vivo* are considerably more complex than those represented in former models, involving the interplay between distinct *cis*-acting elements and the extrinsic termination factor Sen1. We propose an integrated model whereby T-tracts and RNA secondary structures function in concert at primary terminators (and possibly other sites) while Sen1 concurs to release polymerases that have escaped “intrinsic” termination.

### S. cerevisiae Sen1 is a fail-safe transcription termination factor for RNAPIII

One of the important findings of this study is the demonstration that Sen1 can directly promote termination of RNAPIII transcription, a conclusion that is supported both by genome-wide data and by *in vitro* transcription termination assays with purified components.

However, multiple lines of evidence support the notion that Sen1 functions to remove polymerases that have escaped primary termination at the very 3’-end of RNAPIII-dependent genes. These include the high-resolution detection of RNAPIII occupancy by CRAC as well as the analysis of the different RNA species produced from model tRNA genes (figures 2, 4, S2 and S4). A mechanistic explanation for our observation that Sen1 cannot operate at primary terminators is provided by our finding that *in vitro* Sen1 function in termination is hindered by RNA secondary structures, which are typically present in RNAPIII transcripts close to the first terminator.

A recent study has reported that one of the two *Schizosaccharomyces pombe* homologues of Sen1, *Sp* Sen1, also interacts with RNAPIII and its deletion leads to global defects in RNAPIII transcription termination (Rivosecchi et al., 2019). However, some important conclusions of this study are substantially different from the ones supported by our results. It was shown that deletion of *Sp* Sen1 leads to a global downstream shift of the RNAPIII occupancy peak at tRNA genes, as determined by ChIP-seq, and a reduction in the levels of mature tRNAs, which we did not observe in *S. cerevisiae*. These findings have been interpreted in support of a model whereby efficient primary termination in *S. pombe* relies on *Sp* Sen1 and would be only partially dependent on intrinsic termination signals. In contrast, in *S. cerevisiae* primary termination mainly depends on *cis*-signals (T-tracts and secondary structures) and Sen1 operates in downstream regions to remove read-through polymerases. Therefore, in *S. cerevisiae* Sen1 rather plays an important genome safeguarding role in preventing inappropriate extension of RNAPIII transcription.

The divergency between these models might be due to the different resolution of the techniques employed in the two studies (e.g. CRAC and ChIP-seq) but might also reflect mechanistic differences between two organisms. For instance, differences in the biochemical properties of the two Sen1 proteins or in the mode they are loaded onto the nascent transcript might be at stake. Indeed, substantial sequence homology between the two proteins can be found only in their helicase domains and, contrary to *S. cerevisiae* Sen1, none of the *S. pombe* Sen1 homologues is essential for viability, interacts with RNAPII or partakes in RNAPII transcription termination (Larochelle et al., 2018; Rivosecchi et al., 2019).

Importantly, despite the possible mechanistic differences, the fact that the two Sen1 orthologues play a role on RNAPIII transcription opens up the possibility that this function is conserved in other organisms.

### The mechanism of Sen1-dependent RNAPIII transcription termination

Although its best-characterized function is the termination of non-coding transcription by RNAPII within the NNS-complex, *S. cerevisiae* Sen1 is also implicated in other processes like the control of R-loop formation, the resolution of transcription-replication conflicts and DNA-repair (Alzu et al., 2012; Appanah et al., 2020; Li et al., 2016; Mischo et al., 2011). The N-terminal domain of Sen1 is an important hub for protein-protein interactions that might modulate these different functions of Sen1 (Appanah et al., 2020; Han et al., 2020). The function of Sen1 in RNAPIII termination *in vivo* strongly depends on the interaction with RNAPIII, which is mediated by a region in the Sen1 N-terminal domain containing the amino acids mutated in Sen1-3 (W773, E774 and W777). This region is not required for termination *in vitro* indicating that it is not a critical molecular determinant of the process of RNAPIII release, and therefore we suggest it drives the recruitment of Sen1 to RNAPIII, which might be a limiting step for termination *in vivo*.

Mutation of the same amino acids in Sen1-3 also prevents the interaction with the replisome components Ctf4 and Mrc1 (Appanah et al., 2020), yet we show that the interactions of Sen1 with the replisome and RNAPIII are not interdependent but rather mutually exclusive. This suggests either that the same surface mediates the interaction with RNAPIII and the replication fork, or that these mutations alter the conformation of two distinct regions of interaction. The observation that Sen1 promotes RNAPIII transcription termination even in the absence of replication (i.e. in G1) indicates that the action of Sen1 is not restricted to situations of transcription-replication conflicts. However, we cannot exclude that such conflicts might trigger Sen1-dependent termination in some circumstances.

Our *in vitro* data strongly support the notion that Sen1 terminates RNAPIII transcription essentially by the same mechanism employed to induce RNAPII release, for which we have previously provided a detailed molecular model (Han et al., 2017; Leonaitė et al., 2017; Porrua and Libri, 2013). We have shown that Sen1 translocates along the nascent RNA and induces a forward motion of paused RNAPII that results in its release from the DNA. The helicase domain of Sen1 retains all the properties that are necessary for transcription termination and we proposed that a particular subdomain protruding from the helicase core (the “prong”) enters the RNA exit channel provoking destabilizing conformational changes in the elongation complex (Leonaitė et al., 2017; discussed in Han and Porrua, 2017). We show here that the helicase domain is also sufficient for RNAPIII transcription termination and that the essential activities involved in translocation (RNA binding and ATP hydrolysis) as well as the “prong” are required. Akin to what was shown for RNAPII, Sen1 “pushes” paused RNAPIII, which either promotes elongation resumption or results in its release from the DNA (figure 6). Whether the outcome of “pushing” (elongation or termination) is determined by alternative, pre-existing conformations of paused RNAPIII or it is stochastic remains to be determined

A former study reported transcription termination defects at RNAPI-dependent genes upon inactivation of Sen1 *in vivo* (Kawauchi et al., 2008). The interpretation of these data is blurred by the fact that Sen1 inhibition can have multiple indirect effects due to its widespread role in termination of RNAPII transcription. However, we have found that Sen1 associates with RNAPI *in vivo* (table 1) and can also promote the release of paused RNAPI *in vitro* (figure S6). Therefore, altogether these data could point at a common mechanism of transcription termination operating at the three eukaryotic RNAPs and relying on the helicase Sen1.

### RNA structures are enhancers of canonical termination signals

The two fundamental steps in RNAPIII transcription termination are RNAPIII pausing and release from the DNA. The most widely accepted model (Arimbasseri et al., 2013), posits that a stretch of Ts in the non-template DNA strand is sufficient for both pausing and release of RNAPIII. An alternative model was proposed by Nielsen and co-authors (Nielsen et al., 2013), according to which, while T-tracts can promote RNAPIII pausing, an RNA hairpin in the vicinity of the paused RNAPIII is the main determinant for the dissociation of the polymerase from the DNA. These fundamental disparities were attributed to differences in the purity of the RNAPIII preparations employed in the studies supporting these models (Arimbasseri et al., 2014; Nielsen and Zenkin, 2014). Here we use a high-purity preparation of the RNAPIII holoenzyme validated in structural and functional analyses (Hoffmann et al., 2015; Vorländer et al., 2018) to investigate the different mechanisms involved in RNAPIII transcription termination.

We find that the capacity of T-tracts to promote RNAPIII pausing is directly linked to the T-tract length, with T4 terminators supporting very little pausing and T≥9 terminators inducing a complete block of RNAPII elongation (figure 5C-D). Our results show that T6 terminators, which are the most frequently found *in vivo* (figure 4F) are not fully efficient in supporting pausing but can induce RNAPIII release in the absence of any adjacent RNA structure (figure 5C and 7B), indicating that RNA secondary structures are not always required for termination. In contrast, in the case of T4 terminators, which are essentially non-functional *in vitro*, we find that an adjacent RNA secondary structure can convert these sequences into moderately efficient terminators (figure 7A). This behaviour likely phenocopies the situation *in vivo* where the tRNA acceptor stem typically folds very close to the primary terminator and might explain why T4 terminators can be found as primary terminators (figure 4F). Although some tracts of 4 Ts are separated by only 1-2 nt from downstream T-tracts and, thus, could be part of longer interrupted termination signals, more isolated T4 terminators appear to function independently (figure S7) and likely in concert with native RNA secondary structures.

Strikingly, in our assays, very long T-tracts (T≥9) are defective in promoting RNAPIII release, and these defects are more pronounced as the length of the T-tract increases. More precisely, we observe that a fraction of RNAPIIIs stall at the proximal portion of these long T-tracts after “reading” only the first 3-6 nt of the T-tract and fail to dissociate from the DNA. Our interpretation of these observations is that RNAPIII can recognize the T-tract in the downstream duplex region, either because of its sequence or because of the particular structure T-tracts impose to the DNA helix (Stefl et al., 2004). Such interactions would stabilize the EC, thus compensating for the destabilizing effect of the weak rU-dA hybrid and the interaction with the unpaired thymidines in the non-template strand within the transcription bubble. Interestingly, we find that an RNA hairpin forming in the vicinity of these long T-tracts can promote full release of stalled RNAPIIIs (figure 7C), suggesting that *in vivo* long T-tracts might require the concomitant presence of an adjacent secondary structure to be fully proficient in transcription termination.

We provide mechanistic evidence that RNA hairpins can form within the RNA exit channel of RNAPIII (figure 7G) to promote termination. Consistent with this finding, a recent structural study has provided evidence that an RNA hairpin can fold within the RNA exit channel of a bacterial polymerase, leading to a rearrangement of the EC (Kang et al., 2018). Interestingly, structural comparisons indicate that eukaryotic polymerases can also accommodate such RNA secondary structures within their RNA exit channels (Kang et al., 2018). Indeed, a very recent structural study on human RNAPI has provided evidence for the presence of double-stranded RNA in the RNA exit channel (Misiaszek et al, https://doi.org/10.1101/2021.05.31.446457). Therefore, we propose that, in the case of RNAPIII, the formation of an RNA hairpin can induce destabilizing conformational changes in the RNAPIII that would contribute to the dissociation of the EC. Importantly, unlike Nielsen and co-authors (Nielsen et al., 2013), we find that an RNA hairpin can only promote efficient release of paused RNAPIIIs when a T-tract resides in the polymerase main channel (figure 7G).

While this work was in progress, a study using a reporter system in human cells provided evidence that an RNA hairpin located close to a short T-tract (T4) can enhance RNAPIII transcription termination *in cellulo* (Verosloff et al., 2021), pointing to an evolutionarily conserved role for RNA structures in termination.

Taken together, our results allow proposing a revisited model for autonomous RNAPIII transcription termination that can partially reconcile former contradictory findings. According to our model, T-tracts are strictly required for termination, but adjacent RNA structures are important auxiliary elements when the length of the T-tract falls outside of the optimal range. Thus, the protein-independent mechanism of termination of RNAPIII transcription has more commonalities with the so-called intrinsic termination pathway for bacterial RNAP than previously appreciated.

### Multiple mechanisms partake in RNAPIII transcription termination

We and others have observed that RNAPIIIs read through the primary terminator quite frequently and termination at downstream regions was proposed to rely on secondary canonical terminators (Turowski et al., 2016). Indeed, tRNA read-through regions contain T-tracts that are more frequent in the sense orientation than in the antisense orientation (figure S8A-B), suggesting they are under positive selection. However, long T-tracts (T>5) are scarce in these regions (figure 4I), suggesting that alternative evolutionary routes have been undertaken for ensuring efficient termination.

We have shown that both RNA structures and the helicase Sen1 can complement the function of short termination signals. These two factors act in a mutually exclusive manner because: i) both employ a similar mechanism likely involving a conformational change initiated at the level of the RNA exit channel of RNAPIII; and ii) the presence of secondary structures in the nascent transcript prevents Sen1 loading. Our data support the idea that Sen1 would play a more prominent role in termination at read-through regions than RNA structures. A possible explanation is that Sen1 can function both at weak terminators and at other pausing sites, while RNA structures can only work when located sufficiently close to a T-tract.

We have observed that the transcripts encoded in the ∼250 bp region immediately downstream of the primary terminator have a lower propensity to fold into secondary structures than the genomic average (figure S8C-E). While this could be partially due to the higher frequency of T-tracts in this region, which lowers the GC content, it might also be a consequence of Sen1 involvement in fail-safe termination. We suggest that “repurposing” the RNAPII transcription termination factor Sen1 for terminating RNAPIII might have a lower evolutionary cost than generating the appropriate arrangements of T-tracts and RNA structures in tRNA read-through regions.

These considerations do not exclude the possibility that more than one mechanism operate in secondary termination for the same gene. This is for instance illustrated by the tH(GUG)G2 gene (figure S2B), where a secondary T8 terminator is present 60 bp downstream of the primary terminator. Termination at this site is independent of Sen1 most likely because a strong secondary structure forms immediately before T8. The fraction of RNAPIII that escape termination at this site terminates at downstream sites, in the apparent absence of strong secondary structures, in a Sen1-dependent manner.

Taken together, our findings reveal the existence of multiple mechanisms cooperating to promote termination of RNAPIII transcription. We propose that RNA structures contribute to the efficiency of primary termination in some instances (i.e. genes with suboptimal terminators), thanks to the natural proximity of the tRNA acceptor stem to the first T-tract, whereas Sen1 would preferentially function at downstream regions (figure 7H). Efficient termination is important for the rapid recycling of RNAPIII for new cycles of transcription and, thus, for maintaining robust expression of tRNAs and other RNAPIII-dependent transcripts that are essential to sustain cell proliferation. Also, it is crucial to prevent or to minimize the conflicts with other transcribing polymerases as well as with other DNA-associated machineries.

## METHODS

### Construction of yeast strains and plasmids

All the strains used in this paper are listed in table S6. Tagging of *RPC160* with the HTP-tag was performed with standard procedures (Longtine et al., 1998; Rigaut et al., 1999) using plasmid pDL599. Plasmid pDL995 for expression of recombinant Sen1 in insect cells was constructed using the SLIC (sequence and ligation-independent cloning) method (Li and Elledge, 2012).

### Co-immunoprecipitation (Co-IP)

For immunoprecipitation of proteins expressed under their own promoter cells were grown on YPD medium. For proteins expressed under the control of the *GAL1* promoter (i.e. full-length Sen1, Sen1 *Δ*NTD and the N-terminal domain of Sen1), cells were grown on rich medium containing 20 g/L of galactose instead of glucose as the carbon source. Cultures (typically 250 mL) were grown to OD_600_∼1 and then cells were collected by centrifugation and resuspended in 1.5 mL of lysis buffer (10 mM sodium phosphate pH 7.5, 200 mM sodium acetate, 0.25% NP-40, 2 mM EDTA, 1 mM EGTA, 5% glycerol) containing protease inhibitors. Suspensions were frozen in liquid nitrogen and lysed using a Retsch MM301 Ball Mill (5 cycles of 3 minutes at 15 Hz). Lysates were clarified by centrifugation at 13 krpm for 30 min at 4°C and, unless otherwise indicated, treated with 20 μg/mL RNase A for 20 min at 25°C prior to immunoprecipitation. For HTP-tagged proteins, the extracts were then incubated with 2.5 mg of IgG-coupled M-280 tosylactivated dynabeads (Thermo Fisher) for 2 h at 4°C with rotation. After incubation, beads were washed three times with lysis buffer and once with H_2_O and used directly in mass spectrometry analyses.

For proteins overexpressed from the *GAL1* promoter IgG sepharose (GE HEathcare) was used instead. After washes with lysis buffer, beads were washed with TEV cleavage buffer (10 mM Tris pH 8, 150 mM NaCl, 0.1 % NP-40, 0.5 mM EDTA, 1 mM DTT, 5% glycerol) and proteins were then eluted by cleaving the protein A moiety with the TEV protease in TEV cleavage overnight at 4°C.

### Mass spectrometry analysis and label-free quantification

Analysis of Sen1 and RNAPIII coimmunoprecipitates by mass spectrometry was conducted by the proteomics core facility of the Institut Jacques Monod. Proteins were digested by adding 0.2 µg of trypsin (Promega, Madison, WI, USA) per sample followed by incubation in 25 mM NH_4_HCO_3_ at 37°C overnight. The resulting peptides were desalted using ZipTip μ-C18 Pipette Tips (Pierce Biotechnology, Rockford, IL, USA) and analyzed using an Orbitrap Fusion equipped with an easy spray ion source and coupled to a nano-LC Proxeon 1200 (Thermo Scientific, Waltham, MA, USA). Peptides were loaded with an online pre-concentration method and separated by chromatography using a Pepmap-RSLC C18 column (0.75 x 750 mm, 2 μm, 100 Å) from Thermo Scientific, equilibrated at 50°C and operating at a flow rate of 300 nl/min. Peptides were eluted by a gradient of solvent A (H_2_O, 0.1 % FA) and solvent B (ACN/H_2_O 80/20, 0.1% FA), the column was first equilibrated 5 min with 95 % of A, then B was raised to 28 % in 105 min and to 40% in 15 min. Finally, the column was washed with 95% of B during 20 min and re-equilibrated with 95% of A during 10 min. Peptide masses were analyzed in the Orbitrap cell in full ion scan mode, at a resolution of 120,000, a mass range of *m/z* 350-1550 and an AGC target of 4.10^5^. MS/MS were performed in the top speed 3 s mode. Peptides were selected for fragmentation by Higher-energy C-trap Dissociation (HCD) with a Normalized Collisional Energy of 27% and a dynamic exclusion of 60 seconds. Fragment masses were measured in an Ion trap in the rapid mode, with and an AGC target of 1.10^4^. Monocharged peptides and unassigned charge states were excluded from the MS/MS acquisition. The maximum ion accumulation times were set to 100 ms for MS and 35 ms for MS/MS acquisitions respectively.

Label-free quantification was done on Progenesis QI for Proteomics (Waters, Milford, MA, USA) in Hi-3 mode for protein abundance calculation. MGF peak files from Progenesis were processed by Proteome Discoverer 2.4 with the Mascot search engine. The Swissprot protein database was typically used for interrogation. A maximum of 2 missed cleavages was authorized. Precursor and fragment mass tolerances were set to 7 ppm and 0.5 Da, respectively. The following post-translational modifications were included as variable: Oxidation (M), Phosphorylation (STY). Spectra were filtered using a 1% FDR using the percolator node.

### UV crosslinking and analysis of cDNA (CRAC)

The CRAC protocol used in this study is derived from Granneman et al. (Granneman et al., 2009) with several modifications as previously described (Candelli et al., 2018). Briefly, 2 L of cells expressing an HTP-tagged version of Rpc160 (the largest subunit of RNAPIII) at the endogenous locus were grown at 30°C to OD_600_∼ 0.6 in CSM-TRP medium. Cells were crosslinked for 50 seconds using a W5 UV crosslinking unit (UVO3 Ltd) and harvested by centrifugation. Cell pellets were washed once with ice-cold 1x PBS and resuspended in 2.4 mL of TN150 buffer (50 mM Tris pH 7.8, 150 mM NaCl, 0.1% NP-40 and 5 mM *β*-mercaptoethanol) per gram of cells in the presence of protease inhibitors (Complete™ EDTA-free Protease Inhibitor Cocktail, Roche). Suspensions were flash frozen in droplets and cells subjected to cryogenic grinding using a Ball Mill MM 400 (5 cycles of 3 minutes at 20 Hz). The resulting frozen lysates were thawed on ice and digested with DNase I (165 units per gram of cells) at 25°C for 1h to solubilize chromatin and then clarified by centrifugation at 16 krpm for 30 min at 4°C.

RNA-protein complexes were immobilized on M-280 tosylactivated dynabeads coupled with rabbit IgGs (10 mg of beads per sample), washed with TN1000 buffer (50 mM Tris pH 7.8, 1 M NaCl, 0.1% NP-40 and 5 mM β-mercaptoethanol), and eluted by digestion with the TEV protease. RNAs were subjected to partial degradation to reduce their size by adding with 0.2 U of RNase cocktail (RNace-IT, Agilent) and the reaction was stopped by the addition of guanidine–HCl to a final concentration of 6 M. RNA-protein complexes were then incubated with Ni-NTA sepharose (Qiagen, 100 μl of slurry per sample) overnight at 4°C and extensively washed. Sequencing adaptors were ligated to the RNA molecules as described in the original procedure. RNA-protein complexes were eluted with elution buffer containing 50 mM Tris pH 7.8, 50 mM NaCl, 150 mM imidazole, 0.1% NP-40 and 5 mM β-mercaptoethanol fractionated using a Gel Elution Liquid Fraction Entrapment Electrophoresis (GelFree) system (Expedeon) following manufacturer’s specifications. The fractions containing Rpc160 were treated with 100 μg of proteinase K, and RNAs were purified and reverse-transcribed using reverse transcriptase Superscript IV (Invitrogen).

The cDNAs were amplified by PCR using LA Taq polymerase (Takara), and then, the PCR reactions were treated with 200 U/mL of Exonuclease I (NEB) for 1 h at 37°C. Finally, the DNA was purified using NucleoSpin columns (Macherey-Nagel) and sequenced on a NextSeq 500 Illumina sequencer.

### Synchronization of cells in G1 and analysis by flow cytometry

2L of cells were synchronized in the G1 phase of the cell cycle by adding 4 mg of *α*-factor. To maintain cells in G1, 8 mg and 4 mg of *α*-factor were subsequently added to the culture after 1h and 2h of incubation at 30°C, respectively. Cells were collected and processed 1h after the last addition of *α*-factor.

To analyse the DNA content of synchronized cells, 2 mL of culture were collected at different time points and cells were harvested by centrifugation. Cell pellets were resuspended in 50 mM sodium citrate buffer and treated with RNase A (QIAGEN) for 2 hours at 50°C, followed by proteinase K (Sigma) treatment for 2 hours at 50°C. Cell aggregates were then dissociated by sonication and 40 μL of cell suspension were incubated with 170 μL of 50 mM sodium citrate buffer containing 0.5 μM Sytox Green (Invitrogen). Data were acquired on a MASCQuant Analyzer (Miltenyi Biotec) and analyzed with FlowJo Software.

### Dataset processing

CRAC reads were demultiplexed using the pyBarcodeFilter script from the pyCRACutility suite (Webb et al., 2014). Next, the 5ʹ adaptor was clipped with Cutadapt and the resulting insert quality-trimmed from the 3ʹ end using Trimmomatic rolling mean clipping (Bolger et al., 2014). We used the pyCRAC script pyFastqDuplicateRemover to collapse PCR duplicates using a 6- nucleotide random tag included in the 3ʹ adaptor. The resulting sequences were reverse complemented with the Fastx reverse complement that is part of the fastx toolkit (http://hannonlab.cshl.edu/fastx_toolkit/) and mapped to the R64 genome with bowtie2 using “-N 1” option. Reads shorter than 20 nt were filtered out after mapping and coverage files were generated and normalized to counts per million (CPM) using the *bamCoverage* tool from the deepTools package (Ramírez et al., 2016) using a bin size of 1.

### Bioinformatic analyses

All sequence files and annotations were obtained from Saccharomyces Genome Database (*S. cerevisiae* genome version R64-2-1). T-tracts were annotated by first searching for sequences containing at least 4 consecutive thymines (for the plus strand) or adenines (for the minus strand) using the unix command line tool *grep* and then generating coordinate files by the *awk* command. The resulting files were then combined into a single BED file (table S7) using BEDOPS suite (Neph et al., 2012) with the “*everything”* option. For each tRNA gene, the primary terminator was defined as the 1^st^ T-tract after the 3’ end of the mature tRNA. Such primary terminators were identified by comparing the mentioned T-tract annotations and the tRNAs annotations with the *closest* tool from BEDTools (Quinlan and Hall, 2010). T-tracts falling within the 700 bp region immediately downstream of the primary terminator of each tRNA gene were identified with the BEDTools *intersect* tool and defined as secondary terminators (table S8).

Reads mapped to different classes of RNAs were summarized by BEDTools *coverage*. Metagene analyses of RNAPIII occupancy were performed with deepTools suite (Ramírez et al., 2016). Strand-specific coverage bigwig files and modified tRNA coordinates (from the 5’ end to the end of 1^st^ T-tract) were used as inputs for the *computeMatrix* tool using a bin size of 1 and the scale-regions mode. The matrices generated for each strand were subsequently combined by the *computeMatrixOperations* tool with the *rbind* option and used as inputs for the *plotProfile* tool to create a summary plot. For heatmap analyses the log2 ratio of the RNAPIII signal in the *sen1-3* mutant relative to the WT was calculated by the *bigwigCompare* tool using a bin size of 1 and the corresponding bigwig coverage files as inputs. Matrices were generated and combined as for metagene analyses and the final matrix was used as the input for the *plotHeatmap* tool. To analyze the correlation between two replicates, the average RNAPIII signal over regions comprising tRNA genes and 500 bp upstream and downstream regions was computed using the *multiBigwigSummary* tool. The resulting tables were used as inputs for the *plotCorrelation* tool to generate scatter plots and calculate the correlation coefficients using the Spearman method.

To annotate tRNA genes read-through regions in the WT and the *sen1-3* mutant, we first determined a threshold below which CRAC signal was considered as background signal. To do so, genomic regions corresponding to protein-coding genes, which are transcribed by RNAPII, were divided into 20 bp non-overlapping windows and the total signal was computed for each of them using normalized coverage files. The value corresponding to the 95% quantile (value below which 95% of windows values fall) was set as threshold. For each tRNA gene, the 1 kb region immediately downstream of the primary terminator was then divided into 20 bp windows with 1 bp overlap and the RNAPIII CRAC signal was computed for all of them. Contiguous windows with values above the threshold were merged and, most often, this resulted in a single read-through region for each gene. When this was not the case, we manually merged the fragmented regions that were separated by small gaps. Final annotations are provided as BED files in table S9.

The efficiency of transcription termination in the WT and the *sen1-3* mutant was estimated by calculating the read-through index defined as the percentage of RNAPIII signal over the read-through regions relative to the signal over tRNA gene regions. The total RNAPIII signal at each region was computed with the UCSC *bigWigAverageOverBed* package (http://genome.ucsc.edu) using the aforementioned annotations.

Data representation and statistical analyses were performed with R using the *ggplot2* and *plyr* (https://cran.r-project.org/web/packages/plyr/index.html) packages.

### RNA analyses

Yeast cells were grown on 30 mL of YPD medium containing the appropriate additives, depending on the experiment, at 30°C to OD_600_ 0.3 to 0.6. Cells were harvested by centrifugation and RNAs were prepared using standard methods. 4 μg of total RNA were reverse-transcribed by the M-MLV reverse transcriptase (New England BioLabs) following the manufacturer specifications and using oligo d(T) and a mixture of random hexamers at 37°C for 45 min. The resulting cDNAs were analysed by quantitative PCR using the LightCycler 480 SYBR Green I Master reagent (Roche) and LightCycler 480 instrument (Roche) using primers specific for the regions to detect (table S5).

For northern blot assays, typically 10 μg of total RNA were loaded onto a 10% denaturing polyacrylamide gel and separated by electrophoresis at 20 W for 2 h. RNAs were then transferred to a nitrocellulose membrane (GE Amersham Hybond^TM^-N^+^) using a wet transfer system (Trans-Blot cell, Bio-Rad) at 100 V for 2 h at 4°C. Membranes were UV cross-linked and hybridized with the corresponding radioactively labeled probe in commercial buffer (Ultrahyb, Ambion) overnight. For abundant RNA species we employed 5’ end labeled DNA oligonucleotides as probes and hybridizations and subsequent washes were performed at 42°C. For RNA species that were very poorly detected using DNA oligonucleotide probes, we employed RNA probes generated by *in vitro* transcription in the presence of *α*^32^P-UTP using the MAXIscript Kit (Ambion). Hybridization was then performed at 68°C overnight, and the membrane was washed two times for 15 min at 42°C with buffer 2x SSC (30 mM sodium citrate pH 7.0, 300 mM NaCl) containing 0.5% SDS and two times for 15 min at 60°C with buffer 0.1x SSC containing 0.1% SDS. After washes, blots were exposed on a phosphorimager screen and finally scanned using a Typhoon scanner (GE healthcare). Images were analysed using the ImageQuant software (GE healthcare).

### Protein purification

RNAPIII and RNAPI were purified from *Saccharomyces cerevisiae* by heparin chromatography, followed by IgG-affinity chromatography and finally anion-exchange using a previously described procedure (Moreno-Morcillo et al., 2014) with the following modifications: for cells lysis and equilibration of the heparin column (GE Healthcare), we used instead a buffer containing 250 mM Tris–HCl pH 8, 250 mM ammonium sulfate, 20% glycerol, 1 mM EDTA, 10 mM MgCl_2_,10 μM ZnCl_2_, 12 mM *β*-mercaptoethanol and a protease inhibitor cocktail (0.3 μg/mL leupeptin, 1.4 μg/mL pepstatin A, 170 μg/mL PMSF and 330 μg/mL benzamidin). Purified RNAPIII and RNAPI were buffer-exchanged to storage buffer (15 mM HEPES pH 7.5, 150 mM ammonium sulfate, 10 mM DTT), concentrated to 14.9 mg/mL (RNAPIII) and 10.4 mg/mL (RNAPI), snap-frozen in liquid nitrogen and stored at -80°C.

Full-length Sen1 was overexpressed from pFL vector in insect cells (*Trichoplusia ni)* using an optimized synthetic gene (GeneArt, Life Technologies) and the baculovirus expression system (Berger et al., 2004). Briefly, Hi5 cells (Thermo Fisher Scientific) expressing a C-terminally His_6_- tagged version of Sen1 were harvested by centrifugation and lysed by sonication at 4°C in ice-cold buffer A1 (50 mM HEPES-NaOH, pH 7.75, 600 mM NaCl, 15% (v/v) glycerol, 5 mM *β*- mercaptoethanol, 10 mM imidazole, 2 mM MgCl_2_) supplemented with a cocktail of protease inhibitors EDTA free (Roche). The lysate was clarified by centrifugation at 15000 rpm for 1h at 4°C, filtered through a 0.45 μm filter and loaded on a 5 mL Protino Ni-NTA agarose prepacked column (Marchery Nagel), equilibrated with buffer A1. To get rid of nucleic acids bound to Sen1, the column was washed with high salt buffer B2 (50 mM HEPES-NaOH, pH 7.75, 1M NaCl, 15% (v/v) glycerol, 5 mM *β*-mercaptoethanol, 2 mM MgCl_2_) and equilibrated back to buffer A1.. Proteins were eluted using a linear gradient from buffer A1 to B (50 mM HEPES-NaOH, pH 7.75, 200 mM NaCl, 10 % (v/v) glycerol, 5 mM *β*-mercaptoethanol, 400 mM imidazole, 2 mM MgCl_2_), diluted with 2 volumes of buffer D1 (50 mM HEPES-NaOH, pH 7.75, 50 mM NaCl, 15 % (v/v) glycerol, 2 mM *β*-mercaptoethanol, 2 mM MgCl_2_) and then loaded onto two tandem 5 mL heparin HP pre-packed columns (GE Healthcare). Sen1 was eluted using a linear gradient from 20% to 100% of buffer B2 containing 50 mM HEPES-NaOH, pH 7.75, 1 M NaCl, 5% (v/v) glycerol, 2 mM MgCl_2_ and 2 mM DTT. Peak fractions were pooled and subjected to size exclusion chromatography using a Superdex 200 16/600 column (GE healthcare) equilibrated in buffer A3 (50 mM HEPES-NaOH, pH 7.75, 300 mM NaCl, 5% (v/v) glycerol, 2 mM Mg acetate, 2 mM DTT). Finally, the fractions of interest were concentrated using an Amicon Ultra-100 centrifugal filter (Millipore), aliquoted, flash frozen in liquid nitrogen and stored at -80°C.

### *In vitro* transcription termination assays

RNAPIII transcription termination assays were performed using essentially the previously described method for RNAPII (Porrua and Libri, 2015b) with some modifications. For each reaction, elongation complexes (ECs) were assembled by annealing 2.5 pmol of 5’-end radioactively labeled RNA primer with 2.5 pmol of template DNA oligo in hybridization buffer (HB) buffer (20 mM Hepes pH 7.6, 100 mM NaCl, 12 mM MgCl_2_, 10 mM DTT). Subsequently, the RNA:DNA hybrids were incubated with 2 pmol of highly purified RNAPIII in transcription reaction buffer (TRB) buffer (20 mM Hepes pH 7.6, 60 mM (NH_4_)_2_SO_4_, 10 mM MgCl_2_, 10% glycerol, 10 mM DTT) at 20°C for 10 min at 550 rpm. Next, 5 pmol of 5’-end biotinylated non-template DNA were added to the mixture and incubated at 20°C for 10 min with shaking. The resulting ternary ECs were mixed with streptavidin beads (Dynabeads MyOne Streptavidin T1 from Invitrogen, 10 μL of slurry per reaction) pre-washed 4 times with TRB buffer containing 0.1% triton X-100 and then incubated at 20°C for 30 min with gentle shaking. After binding, the beads were washed with 1 volume of TRB containing 0.1% triton X-100, then with 1 volume of TRB containing 250 mM (NH_4_)_2_SO_4_, and finally with 1 volume of TRB. After washes, beads were resuspended in 13 μL of TRB buffer. The reaction was started by adding 7 μL of nucleotides mixture (1 mM each in TRB buffer) and incubating at 28°C for 10 min, and then stopped by the addition of 1 μL of 0.5 M EDTA. Beads and supernatant fractions were then collected separately. RNAs in the supernatant were ethanol-precipitated and resuspended in 10 μL of loading buffer containing 1x TBE and 8 M urea and incubated at 95°C for 3 min before loading onto a 10% denaturing polyacrylamide gel. To isolate RNAs from beads, 10 μL of loading buffer was added to the beads and boiled at 95°C for 3 min and then recovered supernatants as “bead fractions”. Finally, sample were subjected to 10% denaturing PAGE, running for 1 h at 40 W in 1x TBE buffer. Gels were exposed on a phosphorimager screen overnight at -80°C and screens were scanned using a Typhoon scanner (GE healthcare). Images were analysed using the ImageQuant software (GE healthcare).

## DATA AVAILABILITY

The RNAPIII CRAC data have been deposited in NCBI’s Gene Expression Omnibus (GEO) and are accessible through GEO Series accession number GSE174738.

## ACKNOWLEDGEMENTS

We thank G. Wentzinger for technical assistance and other members of the Libri lab for fruitful discussions. We thank F. Fiorini and Hervé Le Hir for sharing the Upf1 protein. We thank the Roscoff Bioinformatics platform ABiMS (http://abims.sb-roscoff.fr) for providing computational resources and support. This work has benefited from the facilities and expertise of the high throughput sequencing core facility of I2BC (http://www.i2bc.paris-saclay.fr/). We thank the proteomics facility of the Institut Jacques Monod, supported by the Region Ile-de-France (SESAME), Université de Paris and the CNRS, for their technical assistance.

## AUTHORS CONTRIBUTIONS

J.X. conducted all experiments except for those in table 1, performed bioinformatic analyses together with Y.C. and contributed to manuscript writing. U.A. and N.H. performed experiments in table 1. M.G. and C.W.M. purified RNAPIII and RNAPI and provide experimental advice. J.S. and V.P. purified Sen1 protein. O.P. conceived the project. O.P. and D.L. designed experiments, supervised research and wrote the manuscript with inputs from all coauthors.

## REFERENCES

Alzu, A., Bermejo, R., Begnis, M., Lucca, C., Piccini, D., Carotenuto, W., Saponaro, M., Brambati, A., Cocito, A., Foiani, M., et al. (2012). Senataxin associates with replication forks to protect fork integrity across RNA-polymerase-II-transcribed genes. Cell 151, 835–846.

Appanah, R., Lones, E.C., Aiello, U., Libri, D., and De Piccoli, G. (2020). Sen1 Is Recruited to Replication Forks via Ctf4 and Mrc1 and Promotes Genome Stability. Cell Rep 30, 2094–2105.e9.

Arimbasseri, A.G., and Maraia, R.J. (2015). Mechanism of Transcription Termination by RNA Polymerase III Utilizes a Non-template Strand Sequence-Specific Signal Element. Mol. Cell 58, 1124–1132.

Arimbasseri, A.G., Rijal, K., and Maraia, R.J. (2013). Transcription termination by the eukaryotic RNA polymerase III. Biochim. Biophys. Acta 1829, 318–330.

Arimbasseri, A.G., Kassavetis, G.A., and Maraia, R.J. (2014). Transcription. Comment on “Mechanism of eukaryotic RNA polymerase III transcription termination.” Science 345, 524.

Arndt, K.M., and Reines, D. (2015). Termination of Transcription of Short Noncoding RNAs by RNA Polymerase II. Annual Review of Biochemistry 84, 381–404.

Baejen, C., Andreani, J., Torkler, P., Battaglia, S., Schwalb, B., Lidschreiber, M., Maier, K.C., Boltendahl, A., Rus, P., Esslinger, S., et al. (2017). Genome-wide Analysis of RNA Polymerase II Termination at Protein-Coding Genes. Molecular Cell 66, 38–49.e6.

Berger, I., Fitzgerald, D.J., and Richmond, T.J. (2004). Baculovirus expression system for heterologous multiprotein complexes. Nat Biotechnol 22, 1583–1587.

Bolger, A.M., Lohse, M., and Usadel, B. (2014). Trimmomatic: a flexible trimmer for Illumina sequence data. Bioinformatics 30, 2114–2120.

Bridier-Nahmias, A., Tchalikian-Cosson, A., Baller, J.A., Menouni, R., Fayol, H., Flores, A., Saïb, A., Werner, M., Voytas, D.F., and Lesage, P. (2015). Retrotransposons. An RNA polymerase III subunit determines sites of retrotransposon integration. Science 348, 585–588.

Candelli, T., Challal, D., Briand, J.-B., Boulay, J., Porrua, O., Colin, J., and Libri, D. (2018). High-resolution transcription maps reveal the widespread impact of roadblock termination in yeast. EMBO J. 37.

Chakrabarti, S., Jayachandran, U., Bonneau, F., Fiorini, F., Basquin, C., Domcke, S., Le Hir, H., and Conti, E. (2011). Molecular mechanisms for the RNA-dependent ATPase activity of Upf1 and its regulation by Upf2. Mol. Cell 41, 693–703.

El Hage, A., Koper, M., Kufel, J., and Tollervey, D. (2008). Efficient termination of transcription by RNA polymerase I requires the 5’ exonuclease Rat1 in yeast. Genes Dev. 22, 1069–1081.

Granneman, S., Kudla, G., Petfalski, E., and Tollervey, D. (2009). Identification of protein binding sites on U3 snoRNA and pre-rRNA by UV cross-linking and high-throughput analysis of cDNAs. Proc. Natl. Acad. Sci. U.S.A. 106, 9613–9618.

Han, Z., and Porrua, O. (2017). Helicases as transcription termination factors: Different solutions for a common problem. Transcription 1–7.

Han, Z., Libri, D., and Porrua, O. (2017). Biochemical characterization of the helicase Sen1 provides new insights into the mechanisms of non-coding transcription termination. Nucleic Acids Res. 45, 1355–1370.

Han, Z., Jasnovidova, O., Haidara, N., Tudek, A., Kubicek, K., Libri, D., Stefl, R., and Porrua, O. (2020). Termination of non-coding transcription in yeast relies on both an RNA Pol II CTD interaction domain and a CTD-mimicking region in Sen1. EMBO J. e101548.

Hazelbaker, D.Z., Marquardt, S., Wlotzka, W., and Buratowski, S. (2013). Kinetic competition between RNA Polymerase II and Sen1-dependent transcription termination. Mol. Cell 49, 55– 66.

Hoffmann, N.A., Jakobi, A.J., Moreno-Morcillo, M., Glatt, S., Kosinski, J., Hagen, W.J.H., Sachse, C., and Müller, C.W. (2015). Molecular structures of unbound and transcribing RNA polymerase III. Nature 528, 231–236.

Kang, J.Y., Mishanina, T.V., Bellecourt, M.J., Mooney, R.A., Darst, S.A., and Landick, R. (2018). RNA Polymerase Accommodates a Pause RNA Hairpin by Global Conformational Rearrangements that Prolong Pausing. Molecular Cell 69, 802–815.e1.

Kawauchi, J., Mischo, H., Braglia, P., Rondon, A., and Proudfoot, N.J. (2008). Budding yeast RNA polymerases I and II employ parallel mechanisms of transcriptional termination. Genes Dev. 22, 1082–1092.

Kim, M., Krogan, N.J., Vasiljeva, L., Rando, O.J., Nedea, E., Greenblatt, J.F., and Buratowski, S. (2004). The yeast Rat1 exonuclease promotes transcription termination by RNA polymerase II. Nature 432, 517–522.

Kireeva, M.L., Komissarova, N., Waugh, D.S., and Kashlev, M. (2000). The 8-nucleotide-long RNA:DNA hybrid is a primary stability determinant of the RNA polymerase II elongation complex. J. Biol. Chem. 275, 6530–6536.

Landrieux, E., Alic, N., Ducrot, C., Acker, J., Riva, M., and Carles, C. (2006). A subcomplex of RNA polymerase III subunits involved in transcription termination and reinitiation. EMBO J 25, 118–128.

Larochelle, M., Robert, M.-A., Hébert, J.-N., Liu, X., Matteau, D., Rodrigue, S., Tian, B., Jacques, P.-É., and Bachand, F. (2018). Common mechanism of transcription termination at coding and noncoding RNA genes in fission yeast. Nat Commun 9.

Leonaitė, B., Han, Z., Basquin, J., Bonneau, F., Libri, D., Porrua, O., and Conti, E. (2017). Sen1 has unique structural features grafted on the architecture of the Upf1-like helicase family. EMBO J. 36, 1590–1604.

Li, M.Z., and Elledge, S.J. (2012). SLIC: a method for sequence- and ligation-independent cloning. Methods Mol Biol 852, 51–59.

Li, W., Selvam, K., Rahman, S.A., and Li, S. (2016). Sen1, the yeast homolog of human senataxin, plays a more direct role than Rad26 in transcription coupled DNA repair. Nucleic Acids Res. 44, 6794–6802.

Longtine, M.S., McKenzie, A., 3rd, Demarini, D.J., Shah, N.G., Wach, A., Brachat, A., Philippsen, P., and Pringle, J.R. (1998). Additional modules for versatile and economical PCR-based gene deletion and modification in Saccharomyces cerevisiae. Yeast 14, 953–961.

Merkl, P., Perez-Fernandez, J., Pilsl, M., Reiter, A., Williams, L., Gerber, J., Böhm, M., Deutzmann, R., Griesenbeck, J., Milkereit, P., et al. (2014). Binding of the termination factor Nsi1 to its cognate DNA site is sufficient to terminate RNA polymerase I transcription in vitro and to induce termination in vivo. Mol. Cell. Biol. 34, 3817–3827.

Mischo, H.E., Gomez-Gonzalez, B., Grzechnik, P., Rondon, A.G., Wei, W., Steinmetz, L., Aguilera, A., and Proudfoot, N.J. (2011). Yeast Sen1 helicase protects the genome from transcription-associated instability. Mol Cell 41, 21–32.

Mishra, S., and Maraia, R.J. (2019). RNA polymerase III subunits C37/53 modulate rU:dA hybrid 3’ end dynamics during transcription termination. Nucleic Acids Res 47, 310–327.

Moreno-Morcillo, M., Taylor, N.M.I., Gruene, T., Legrand, P., Rashid, U.J., Ruiz, F.M., Steuerwald, U., Müller, C.W., and Fernández-Tornero, C. (2014). Solving the RNA polymerase I structural puzzle. Acta Crystallogr D Biol Crystallogr 70, 2570–2582.

Neph, S., Kuehn, M.S., Reynolds, A.P., Haugen, E., Thurman, R.E., Johnson, A.K., Rynes, E., Maurano, M.T., Vierstra, J., Thomas, S., et al. (2012). BEDOPS: high-performance genomic feature operations. Bioinformatics 28, 1919–1920.

Nielsen, S., and Zenkin, N. (2014). Transcription. Response to Comment on “Mechanism of eukaryotic RNA polymerase III transcription termination.” Science 345, 524.

Nielsen, S., Yuzenkova, Y., and Zenkin, N. (2013). Mechanism of eukaryotic RNA polymerase III transcription termination. Science 340, 1577–1580.

Osmundson, J.S., Kumar, J., Yeung, R., and Smith, D.J. (2017). Pif1-family helicases cooperatively suppress widespread replication-fork arrest at tRNA genes. Nat Struct Mol Biol 24, 162–170.

Park, J., Kang, M., and Kim, M. (2015). Unraveling the mechanistic features of RNA polymerase II termination by the 5ʹ-3ʹ exoribonuclease Rat1. Nucleic Acids Res 43, 2625–2637.

Pearson, E.L., and Moore, C.L. (2013). Dismantling Promoter-driven RNA Polymerase II Transcription Complexes in Vitro by the Termination Factor Rat1. J. Biol. Chem. 288, 19750– 19759.

Porrua, O., and Libri, D. (2013). A bacterial-like mechanism for transcription termination by the Sen1p helicase in budding yeast. Nat. Struct. Mol. Biol. 20, 884–891.

Porrua, O., and Libri, D. (2015a). Transcription termination and the control of the transcriptome: why, where and how to stop. Nat. Rev. Mol. Cell Biol. 16, 190–202.

Porrua, O., and Libri, D. (2015b). Characterization of the mechanisms of transcription termination by the helicase Sen1. Methods Mol. Biol. 1259, 313–331.

Porrua, O., Hobor, F., Boulay, J., Kubicek, K., D’Aubenton-Carafa, Y., Gudipati, R.K., Stefl, R., and Libri, D. (2012). In vivo SELEX reveals novel sequence and structural determinants of Nrd1-Nab3-Sen1-dependent transcription termination. EMBO J. 31, 3935–3948.

Porrua, O., Boudvillain, M., and Libri, D. (2016). Transcription Termination: Variations on Common Themes. Trends Genet. 32, 508–522.

Quinlan, A.R., and Hall, I.M. (2010). BEDTools: a flexible suite of utilities for comparing genomic features. Bioinformatics 26, 841–842.

Ramírez, F., Ryan, D.P., Grüning, B., Bhardwaj, V., Kilpert, F., Richter, A.S., Heyne, S., Dündar, F., and Manke, T. (2016). deepTools2: a next generation web server for deep-sequencing data analysis. Nucleic Acids Res. 44, W160–165.

Reiter, A., Hamperl, S., Seitz, H., Merkl, P., Perez-Fernandez, J., Williams, L., Gerber, J., Németh, A., Léger, I., Gadal, O., et al. (2012). The Reb1-homologue Ydr026c/Nsi1 is required for efficient RNA polymerase I termination in yeast. EMBO J. 31, 3480–3493.

Rigaut, G., Shevchenko, A., Rutz, B., Wilm, M., Mann, M., and Séraphin, B. (1999). A generic protein purification method for protein complex characterization and proteome exploration. Nat. Biotechnol. 17, 1030–1032.

Rijal, K., and Maraia, R.J. (2013). RNA polymerase III mutants in TFIIFα-like C37 that cause terminator readthrough with no decrease in transcription output. Nucleic Acids Res 41, 139– 155.

Rivosecchi, J., Larochelle, M., Teste, C., Grenier, F., Malapert, A., Ricci, E.P., Bernard, P., Bachand, F., and Vanoosthuyse, V. (2019). Senataxin homologue Sen1 is required for efficient termination of RNA polymerase III transcription. EMBO J 38, e101955.

Schulz, D., Schwalb, B., Kiesel, A., Baejen, C., Torkler, P., Gagneur, J., Soeding, J., and Cramer, P. (2013). Transcriptome Surveillance by Selective Termination of Noncoding RNA Synthesis. Cell.

Skowronek, E., Grzechnik, P., Späth, B., Marchfelder, A., and Kufel, J. (2014). tRNA 3’ processing in yeast involves tRNase Z, Rex1, and Rrp6. RNA 20, 115–130.

Stefl, R., Wu, H., Ravindranathan, S., Sklenár, V., and Feigon, J. (2004). DNA A-tract bending in three dimensions: solving the dA4T4 vs. dT4A4 conundrum. Proc Natl Acad Sci U S A 101, 1177–1182.

Steinmetz, E.J., Warren, C.L., Kuehner, J.N., Panbehi, B., Ansari, A.Z., and Brow, D.A. (2006). Genome-wide distribution of yeast RNA polymerase II and its control by Sen1 helicase. Mol. Cell 24, 735–746.

Turowski, T.W., Leśniewska, E., Delan-Forino, C., Sayou, C., Boguta, M., and Tollervey, D. (2016). Global analysis of transcriptionally engaged yeast RNA polymerase III reveals extended tRNA transcripts. Genome Res 26, 933–944.

Verosloff, M.S., Corcoran, W.K., Dolberg, T.B., Bushhouse, D.Z., Leonard, J.N., and Lucks, J.B. (2021). RNA Sequence and Structure Determinants of Pol III Transcriptional Termination in Human Cells. J Mol Biol 433, 166978.

Vorländer, M.K., Khatter, H., Wetzel, R., Hagen, W.J.H., and Müller, C.W. (2018). Molecular mechanism of promoter opening by RNA polymerase III. Nature 553, 295–300.

Wang, S., Han, Z., Libri, D., Porrua, O., and Strick, T.R. (2019). Single-molecule characterization of extrinsic transcription termination by Sen1 helicase. Nat Commun 10, 1545.

Webb, S., Hector, R.D., Kudla, G., and Granneman, S. (2014). PAR-CLIP data indicate that Nrd1-Nab3-dependent transcription termination regulates expression of hundreds of protein coding genes in yeast. Genome Biol 15, R8.

West, S., Gromak, N., and Proudfoot, N.J. (2004). Human 5’ --> 3’ exonuclease Xrn2 promotes transcription termination at co-transcriptional cleavage sites. Nature 432, 522–525.

Wlotzka, W., Kudla, G., Granneman, S., and Tollervey, D. (2011). The nuclear RNA polymerase II surveillance system targets polymerase III transcripts. EMBO J. 30, 1790–1803.

Yüce, Ö., and West, S.C. (2013). Senataxin, defective in the neurodegenerative disorder ataxia with oculomotor apraxia 2, lies at the interface of transcription and the DNA damage response. Mol. Cell. Biol. 33, 406–417.

