## Supplementary material for "An integrated model for termination of RNA polymerase III transcription"

**List of supplementary material:**

- Figures S1-S8.

-Table S6.

**Material provided as separate xls files:**

-Table S1: mass spectrometry analyses of TAP-Sen1 and TAP-Sen1 ΔNTD coimmunoprecipitates.

-Table S2: mass spectrometry analyses of TAP-Sen1 NTD coimmunoprecipitates.

-Table S3: mass spectrometry analyses of Sen1-TAP and Sen1-3-TAP coimmunoprecipitates.

-Table S4: label-free quantitative mass spectrometry analyses of Rpc160-HTP coimmunoprecipitates in a WT and a *sen1-3* background.

-Table S5: list of oligonucleotides used in this study.

-Table S7: annotations of tRNA genes from the 5' end to the mature tRNA to the primary terminator in bed format.

-Table S8: annotations of potential secondary terminators of tRNA genes in bed format.

-Table S9: annotations of tRNA genes readthrough regions in a WT and a *sen1-3* mutant in bed format.

### SUPPLEMENTARY FIGURES

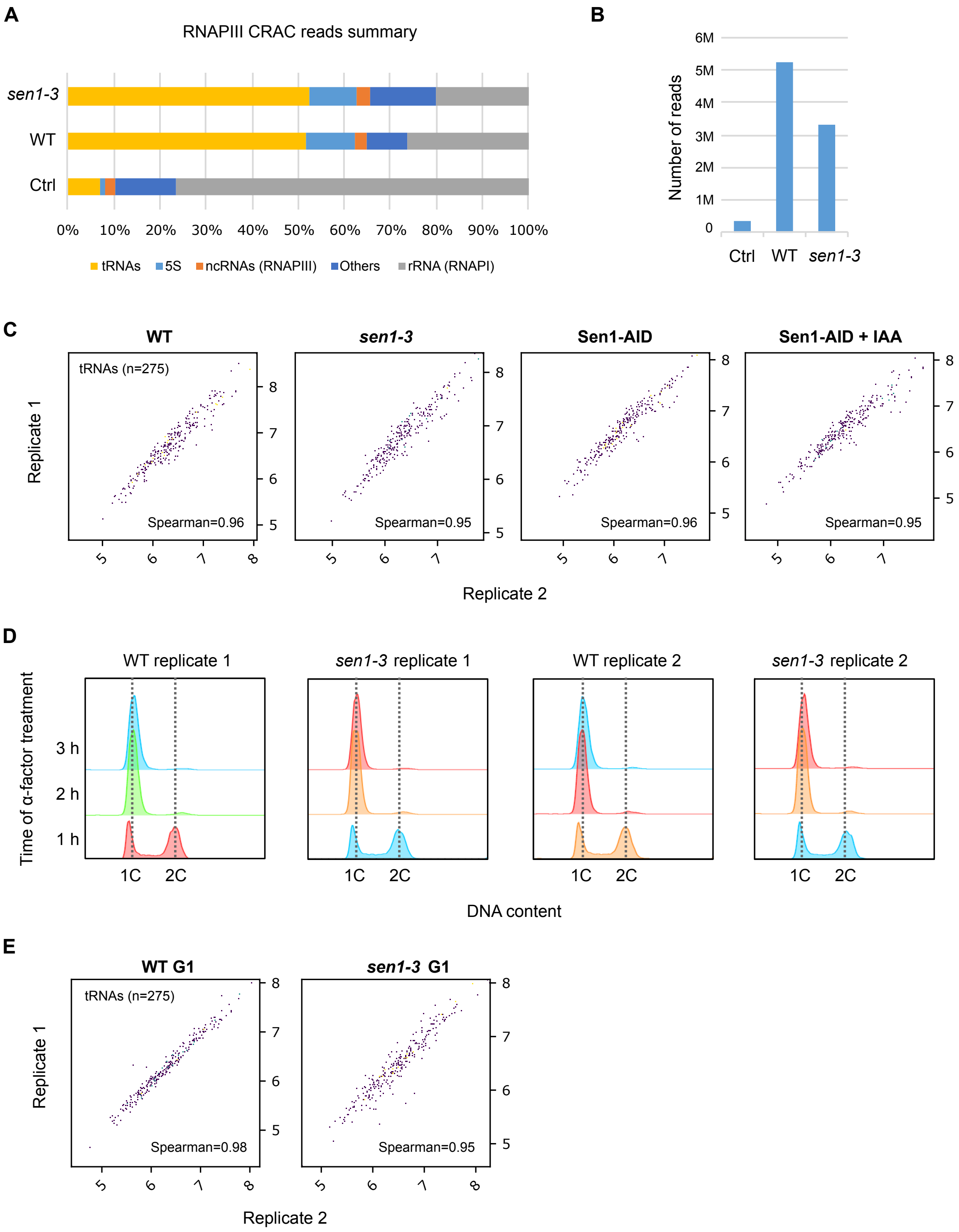


**Figure S1: Complementary analyses validating CRAC experiments in figures 2 and 3.**
**A)** Comparison of the reads distribution among different genomic regions in crosslinked samples (WT and sen1-3) relative to the un-crosslinked control (Ctrl) in a typical RNAPIII CRAC experiment. The "others" category corresponds to RNAPII genes and intergenic regions. Note that tRNA read-through regions are included in this category and the larger proportion of

reads in this group in the *sen1-3* mutant could be due to the observed increased RNAPIII presence at those regions.

**B)** Plot representing the number of mapped reads obtained in a typical CRAC experiment in the different samples. Note that the number of reads in cross-linked samples is at least one order of magnitude higher than in the un-crosslinked control (Ctrl). "M" denotes millions.

**C)** Scatter plots showing the high correlation between the two biological replicates of each condition/strain for the CRAC experiments showed in figure 2.

**D)** Analysis of DNA copy-number for samples in figure 3D-F by flow cytometry. 1C and 2C corresponds to 1 and 2 copies of the genome, respectively. Cultures were used for CRAC analyses after 3h of treatment with $\alpha$-factor (see methods).

**E)** Correlation plots for the two biological replicates of samples used in experiments in figure 3D-F.

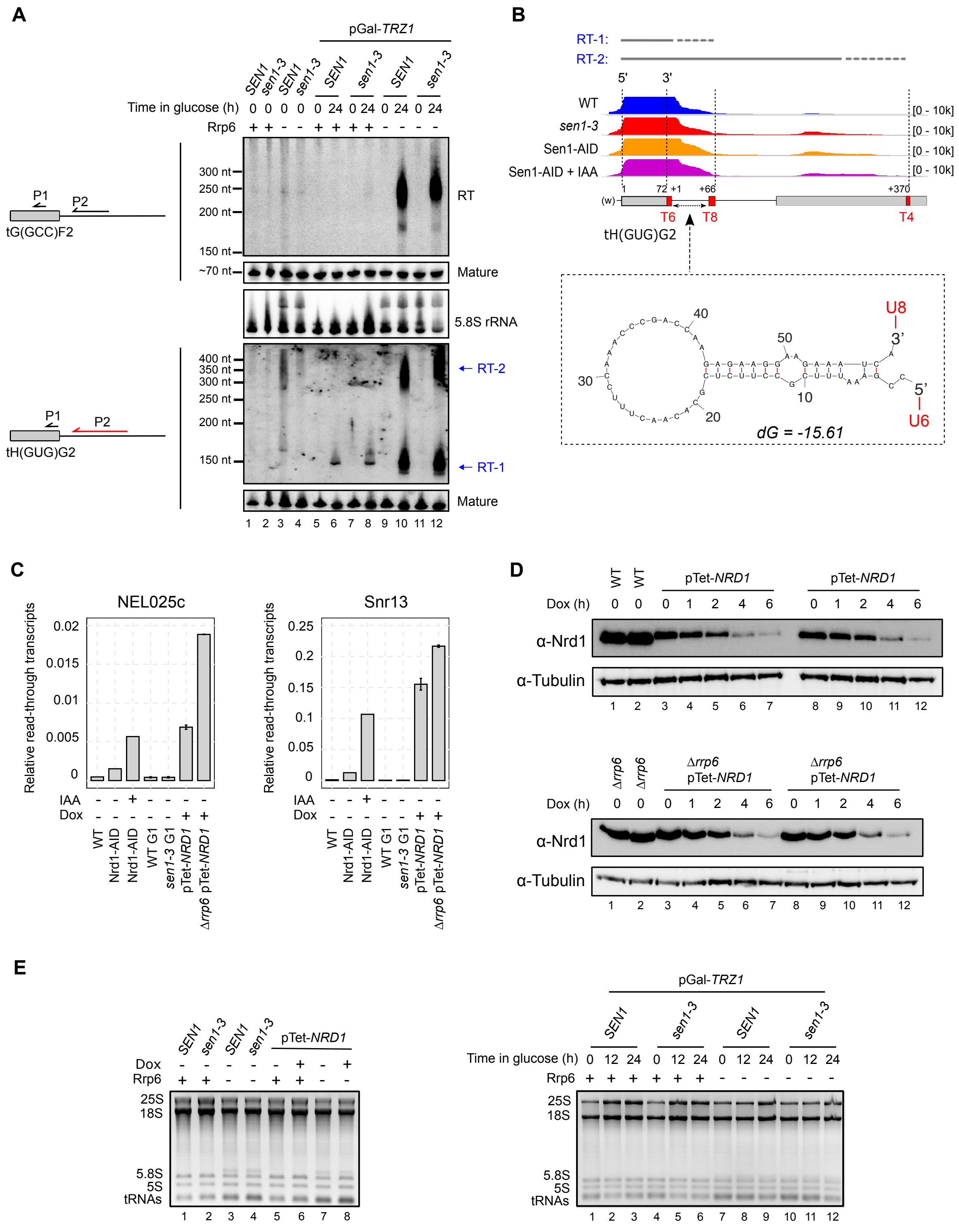


**Figure S2: Experiments related to figures 2 and 3.**

**A)** Northern blot analysis of transcripts derived from two different tRNA genes in the indicated backgrounds. In pGal-*TRZ1* strains, the essential gene *TRZ1* is expressed from the *GAL1* promoter and the different strains are either grown on galactose for the whole experiment (t=0) or shifted to medium containing glucose for 24h to repress *TRZ1.* Experiments performed with 12h-incubation in glucose-containing medium provided similar results (data not shown). Schemes on the left indicate the approximate position of the probes (P1 and P2) used for the detection of the different RNA species (RT, for read-through, and mature tRNA). The RNA probe is indicated in red, while DNA oligonucleotide probes are indicated in black (more details in table S5). The 5.8S rRNA is used as a loading control.

**B)** IGV screenshot of the region around the tH(GUG)G2 gene indicating the position of different T-tracts found and the two major groups of RT transcripts detected by northern blot (**A**, bottom blot). Note that the fact that we detect multiple bands for each termination region could be due to the existence of several termination sites and/or the presence of heterogeneous poly(A) tails. The structure of the RNA between the T6 and the T8 T-tract predicted by the mFold software of the UNAFold package (http://www.unafold.org/) is shown on the bottom.

**C)** Western blot analysis of Nrd1 depletion by incubation of pTet-*NRD1* strains (2 biological replicates) with doxycycline (Dox) for the indicated times. Tubulin detection was used as a loading control.

**D)** Analysis of transcription termination defects at two well-characterized NNS-targets (the CUT NEL025c and the snoRNA gene *SNR13*) in the indicated strains. RNAs where prepared from the same cultures used for CRAC experiments and northern blot analyses in figures 2 and 3. Typical read-through transcripts resulting from inefficient termination by the NNS-complex where detected by RT-qPCR with oligonucleotides listed in table S5. Values are normalized relative to the levels of the *ACT1* mRNA.

**E)** Native agarose gels showing total RNA levels in the indicated samples stained with ethidium bromide.

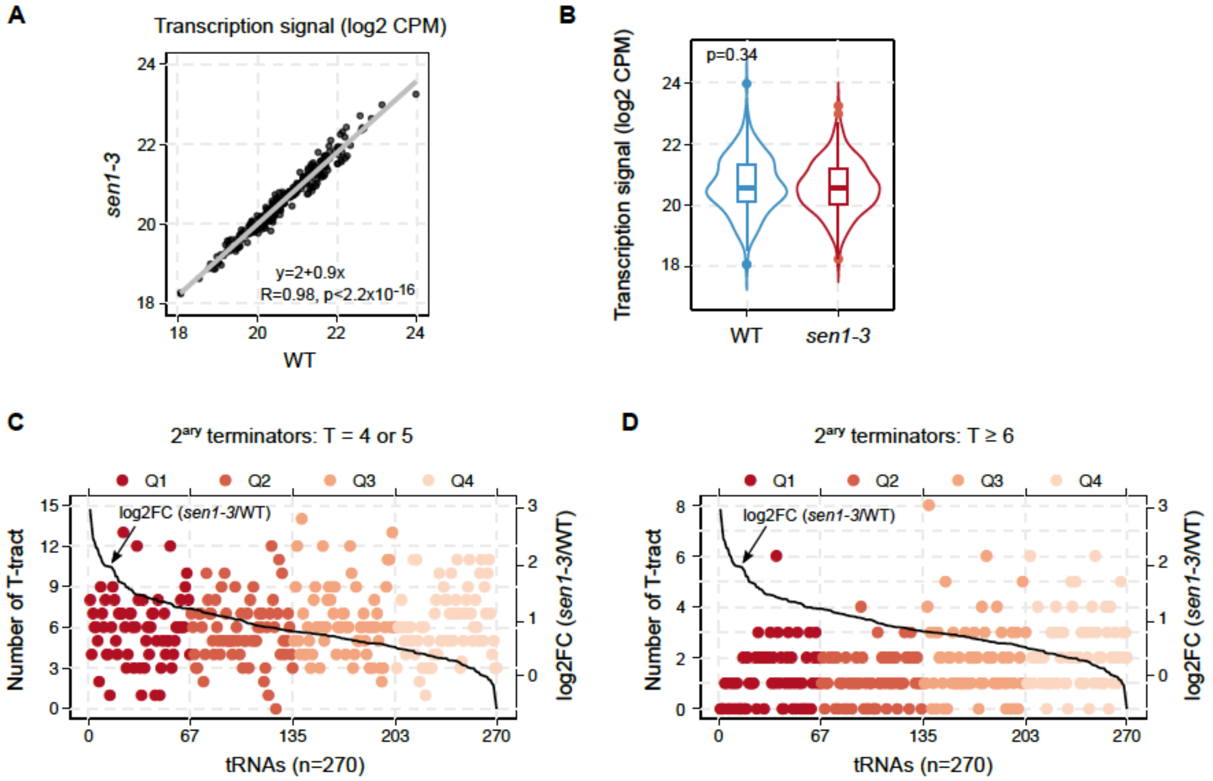


**Figure S3: Complementary analyses related to figure 4.**

**A)** and **B)** Comparison of the transcription signal at tRNA genes in the WT and in the *sen1-3* mutant measured by the total RNAPIII CRAC signal at the region from the 5' end of the mature tRNA to the first T-tract. **A)** Dispersion plot where "R" corresponds to Pearson's correlation coefficient and *p* is the associated p-value. **B)** Violin plot where *p* corresponds to the p-value calculated by the Wilcoxon test.

**C)** and **D)** Representation of the number of T-tracts of indicated lengths located in the 700 bp region downstream of the primary terminator of each tRNA gene. Data points are coloured according to the quartile (Q) they belong to. Quartiles are defined according to the log2 FC of the RNAPIII signal in the *sen1-3* relative to the WT at the same region, which provides an estimation of the dependency on Sen1 for termination. Thus, Q1 includes the most Sen1-dependent tRNA genes.

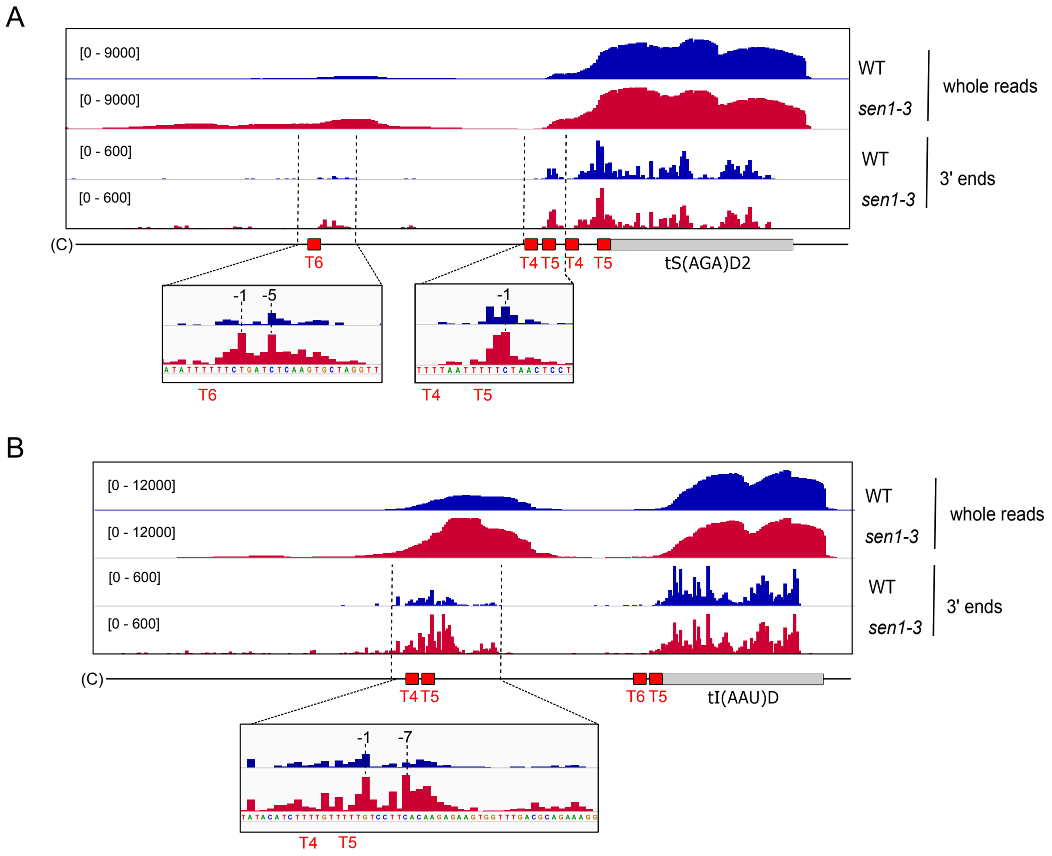


**Figure S4: Examples of tRNA genes illustrating the role of Sen1 in enhancing secondary termination.** IGV screenshots showing the distribution of RNAPIII CRAC signal in the WT and the *sen1-3* mutant with zoom in views of the main regions where RNAPIII accumulates in the mutant. The 3' ends datasets provide the position of individual RNAPIIIs with single-nucleotide resolution. Most accumulation is observed just upstream of or at the first Ts of secondary weak terminators, suggesting impaired RNAPIII release by Sen1 at these sites. The indicated coordinates correspond to the position relative to the beginning of the nearest downstream T-tract.

1) No hairpin and T4, T5, T6, T9 or T12 terminators (figures 5, and 6)

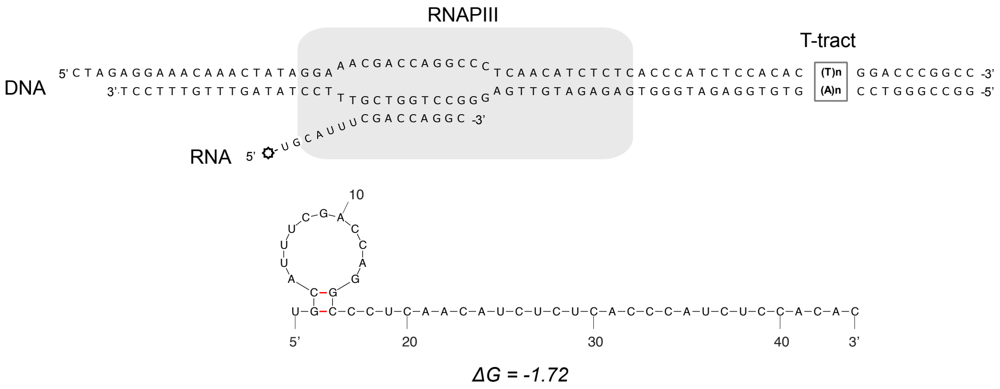


2) Hairpin (HP) and T4, T6 or T12 terminators (figure 7E-F)

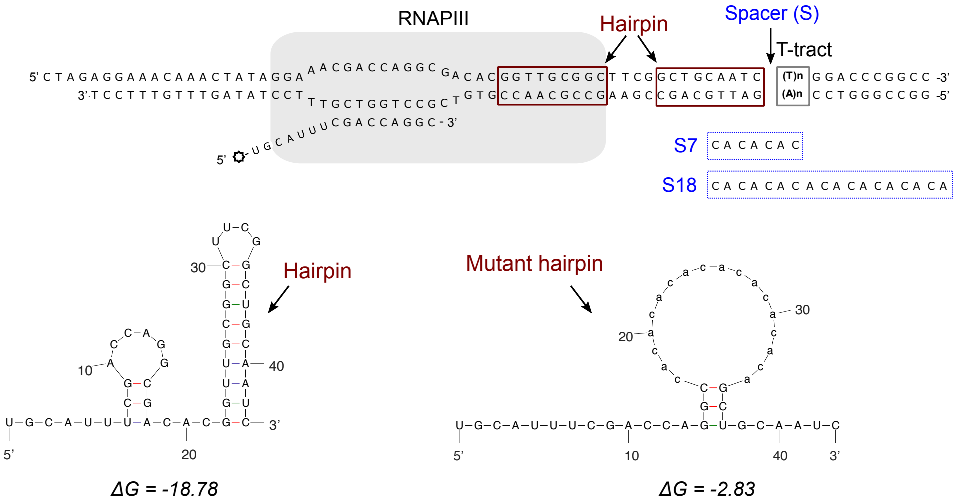


3) Hairpin (HP) - A-less cassette - T4 terminator - A-tract (figure 7E)

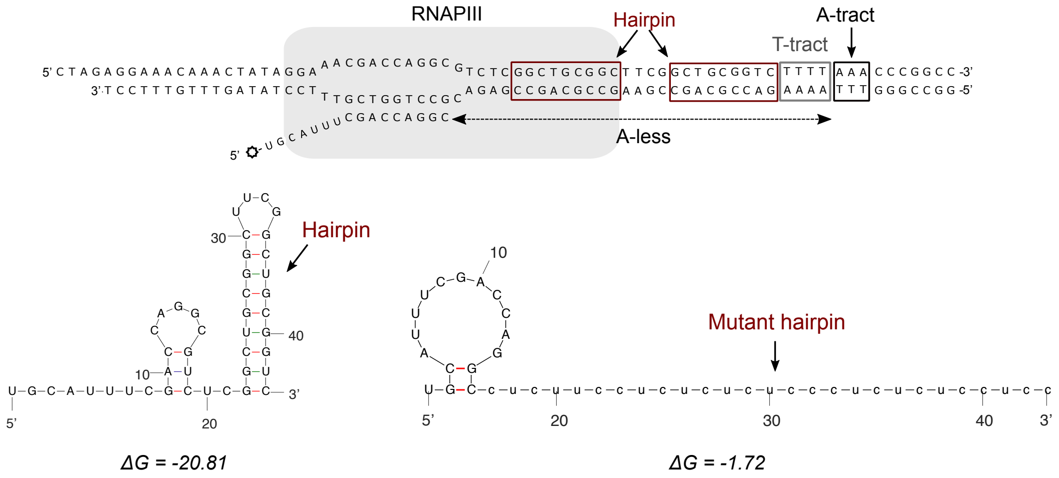


**Figure S5: Sequence of transcription templates and predicted structure of the different transcribed RNAs in *in vitro* transcription termination assays.**

The sequence of the wild-type version of each template are indicated in the schemes. The mutant version of the transcribed RNAs is shown together with the wild-type version under the corresponding scheme. The sequence of the spacers (S) correspond to the non-template strand. RNA structure predictions and ΔG calculation for each structure were performed with the mFold software.

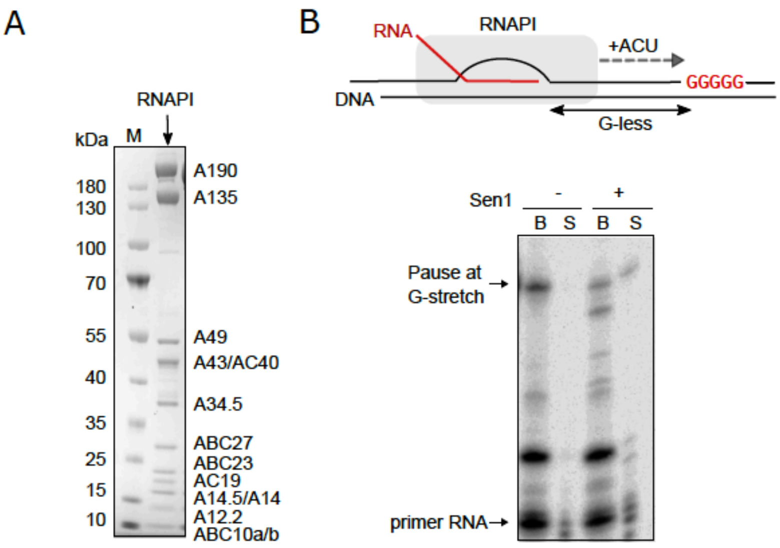


**Figure S6: Sen1 can promote the release of paused RNAPIs *in vitro*.**
**A)** SDS-PAGE analysis of the RNAPI preparation used in these assays.
**B)** IVTT assay performed on templates containing a G-less cassette followed by a run of Gs to promote stalling of RNAPI at the first G in the absence of guanine in the reaction. Top: scheme of the transcription templates. Bottom: Denaturing PAGE analysis of transcripts from one out of two independent IVTT assays, which produced very similar results.

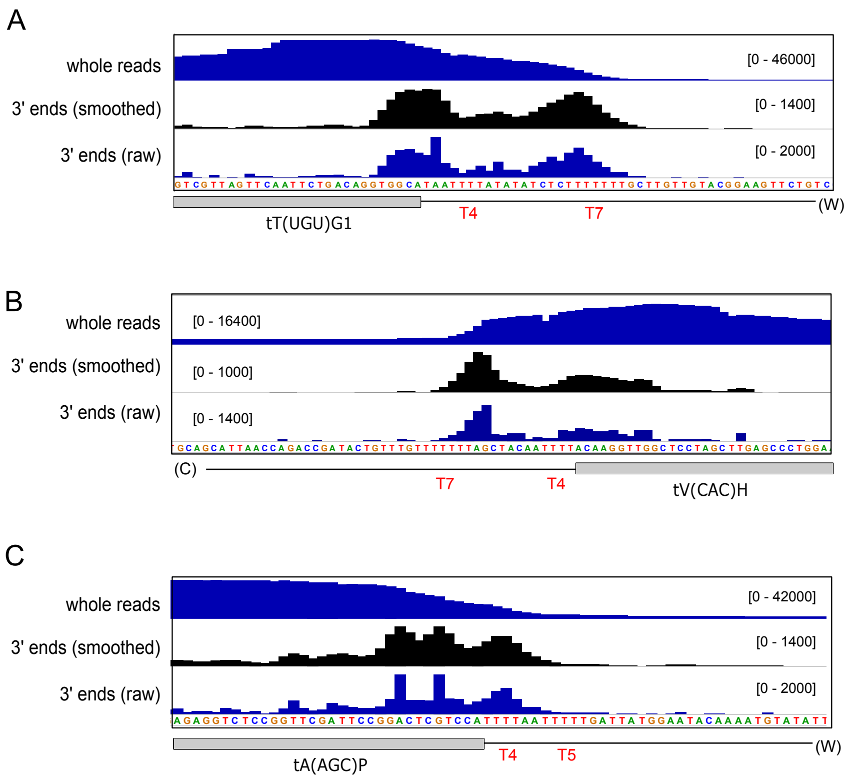


**Figure S7: Examples of tRNA genes harbouring a T4 primary terminator.**

**A)** and **B)** Cases where the T4 terminator seems to function autonomously to promote moderate levels of termination. The 3' end datasets provide accurately the position of individual RNAPIIIs, and therefore the location of pausing sites. The decrease in the whole reads RNAPIII signal within the T4 sequence despite the presence of very strong pausing at close downstream T-tracts supports the idea that a fraction of RNAPIIIs terminate at the T4 terminator.

**C)** Example where the T4 sequence seems to function in combination with the downstream T5 T-tract, as suggested by the strong termination observed at these sequences compared to A) and B), where the T4 sequence is further from other T-tracts. The pausing pattern, with accumulation of RNAPIIIs at the first 3 thymidines of the T4 sequence rather resembles the pattern observed for long T-tracts (e.g. T9 terminators) *in vitro*.

Datasets correspond to the WT strain.

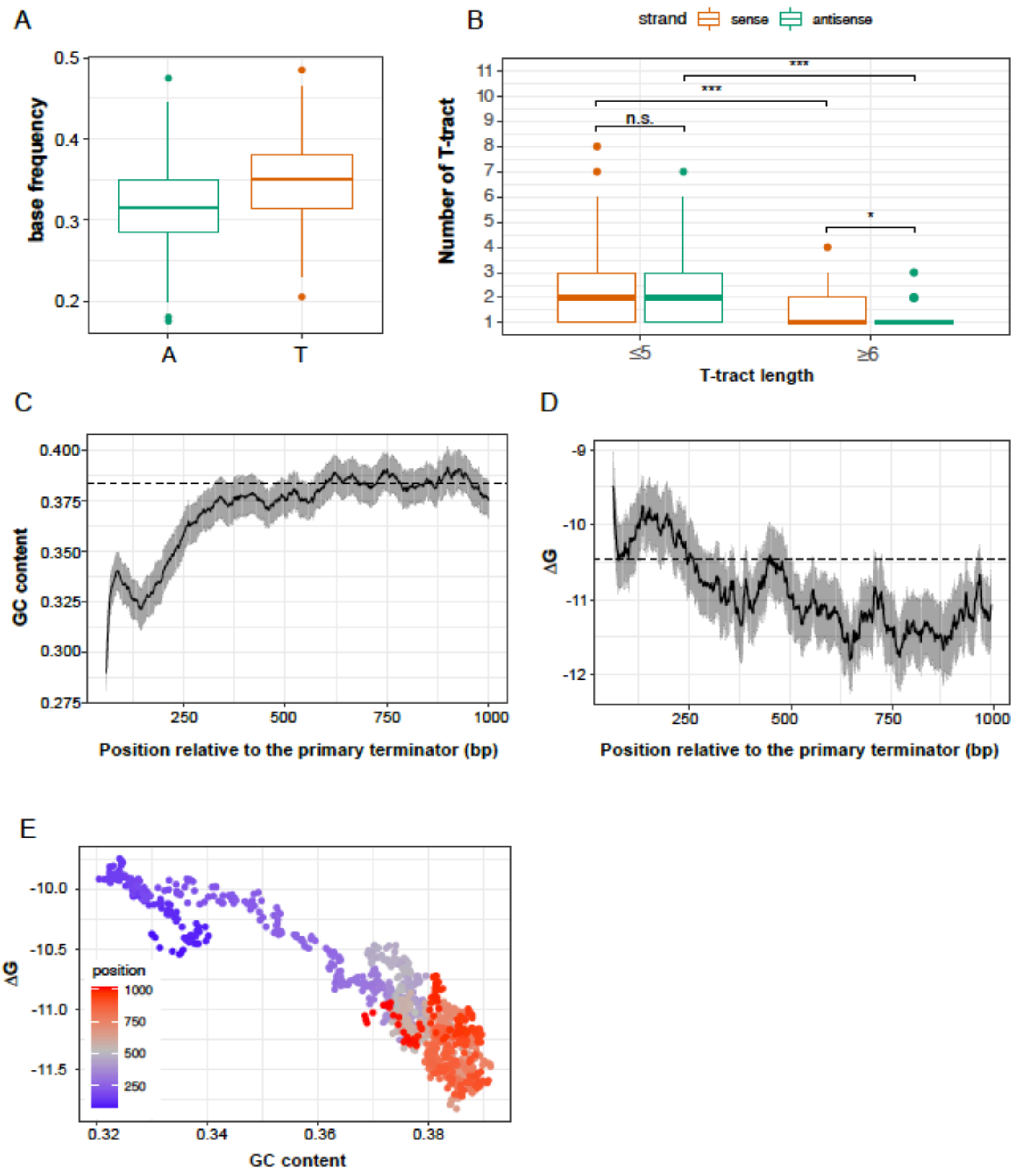


**Figure S8: Analysis of T-tracts and RNA structures at regions of secondary termination.**
**A)** Analysis of the frequency of A and T nucleotides in the 200 bp region downstream of the primary terminator of tRNA genes.
**B)** Comparison of the number of weak (T ≤ 5) or strong (T ≥ 6) terminators at the 200 bp region downstream of the primary terminator of tRNA genes in the sense orientation versus to the antisense orientation relative to transcription. Statistical significance was calculated using a Wilcoxon rank sum test. n.s indicates no significant difference between the compared groups whereas * denotes a p-value ≤ 0.05 and *** a p-value ≤ 0.001.
**C)** Analysis of the GC content (fraction of G and C nucleotides) of regions downstream of the primary terminator of tRNA genes. Values were calculated for 60 bp sliding windows. The black line corresponds to the average value whereas the grey zone represents the 95%

confidence interval of the average value. A dashed line indicates the average value for the whole genome.

**D)** Analysis of the Gibbs free energy ($\Delta G$) as a proxy for the propensity of the transcribed regions to form secondary structures. We plotted values calculated for 65 bp sliding windows published in Turowski et al. 2020. A dashed line indicates the average $\Delta G$ value for the whole genome.

**E)** Combined representation of the GC content and the $\Delta G$ of the different regions coloured according to the distance from the primary terminator. Closer regions (blue) tend to be less GC-rich and less structured while further regions (red) tend to be more GC-rich and structured.

**Table S6**: Yeast strains used in this study.

| **Number** | **Name** | **Genotype** | **Source** |
|---|---|---|---|
| DLY671 | BMA | as W303, Δ*trp1* | F. Lacroute |
| DLY1152 | *Δrrp6* | as W303*, rrp6::URA3* | F. Lacroute |
| DLY1605 | *pTet-NRD1* | as BMA, *pTet::FLAG::NRD1* | J. Colin |
| DLY1626 | *pTet-NRD1, Δrrp6* | as BMA*, pTet::FLAG::NRD1, rrp6::KAN* | This work |
| DLY1656 | $P_{GAL1-}$ *TAP-SEN1* | as BMA, *TRP1::Pgal::TAP::SEN1* | Porrua et al, 2012 |
| DLY2692 | $P_{GAL1-}$ *TAP-sen1ΔNter* | as BMA, *TRP1::Pgal::TAP::sen1ΔNter (Δ1-975)* | Han et al, 2020 |
| DLY3171 | Sen1-TAP | as W303, *SEN1::TAP::KAN* | Appanah et al, 2020 |
| DLY3173 | *sen1-3*-TAP | as W303, *sen1W773A,E774A,W777A::TAP::KAN* | Appanah et al, 2020 |
| DLY3197 | *sen1-3* | as W303, *sen1W773A,E774A,W777A* | This work |
| DLY3246 | *sen1-3, Δrrp6* | *as W303, sen1W773A,E774A,W777A rrp6::URA3* | This work |
| DLY3262 | Rpc160-HTP | as BMA, *RPC160::HTP::TRP1* | This work |
| DLY3265 | Rpc160-HTP, *sen1-3* | as W303, RPC160::HTP::TRP1, *sen1W773A,E774A,W777A* | This work |
| DLY3343 | Rpc160-HTP, Sen1-AID | as BMA, *RPC160::HTP::TRP1, SEN1-AID::KAN::OsTIR1* | This work |
| DLY3377 | Rpc160-HTP, Nrd1-AID | as BMA, *RPC160::HTP::TRP1*, *NRD1-3Flag-AID::KAN::OsTIR1* | This work |
| DLY3462 | GAL::HA-Trz1 | *GAL-HA-Trz1::KAN* | This work |
| DLY3463 | GAL::HA-Trz1, sen1-3 | GAL-HA-Trz1::KAN, *sen1W773A,E774A,W777A* | This work |
| DLY3464 | GAL::HA-Trz1, *Δrrp6* | *GAL-HA-Trz1::KAN, rrp6::URA3* | This work |
| DLY3465 | GAL::HA-Trz1, *Δrrp6,* sen1-3 | *GAL-HA-Trz1::KAN, rrp6::URA3, sen1W773A,E774A,W777A* | This work |
